# The effects of maternal social connectivity and integration on offspring survival in a marmot

**DOI:** 10.1101/2021.01.28.428660

**Authors:** Anita Pilar Montero, Dana M. Williams, Julien G.A. Martin, Daniel T. Blumstein

## Abstract

In social species, maternal social relationships, in addition to direct care, impact offspring survival but much of what we know about these effects comes from studies of obligately social and cooperatively breeding species. Yellow-bellied marmots (*Marmota flaviventer*) are a facultatively social species whose social groups vary in composition, size, and cohesiveness. This natural variation in sociality and cooperative breeding behavior makes yellow-bellied marmots an ideal species within which to study the effects of maternal affiliative and agonistic social behavior on offspring. We used social network analysis to investigate the relationship between maternal social connectivity and integration on offspring summer and yearly survival, with the hypothesis that offspring with more affiliative mothers are more likely to survive than the offspring of more agonistic mothers. However, we found the inverse to be true: pups born to mothers who received more affiliative interactions were less likely to survive while the offspring of mothers who were more highly integrated into agonistic networks had enhanced survival. Overall, maternal social network measures were positively and negatively correlated with offspring survival, indicating that pups are influenced by their mother’s social world, often in contradictory ways. Relative predation risk and colony location also mediated the effects of social relationships on pup survival. This study contributes to a small but growing body of work that demonstrates that specific attributes of sociality have specific consequences and that by adopting an attribute-focused view of sociality we are better able to understand how environmental conditions mediate the costs and benefits of sociality.

**Lay Summary:** Maternal social relationships can impact offspring survival but much of what we know about these effects comes from studies of obligately social species. In faculatively social yellow-bellied marmots we found that pups born to mothers who received more affiliative interactions were less likely to survive while the offspring of mothers who were more highly integrated into agonistic networks had enhanced survival. Overall, pups are influenced by their mother’s social world, often in contradictory ways.

## Introduction

Many taxa exhibit maternal care, where the degree and quality of care directly affects offspring outcomes, particularly their likelihood of survival during the infant and juvenile periods (Diesel 1989, O’Connor and Shine 2004, Grandinson 2005). Maternal care in mammals can take the form of nursing (Stern and Lonstein 2001), grooming (Moore and Power 1992), attentiveness (Lonstein et al. 2015), and protectiveness of offspring towards threatening conspecifics (Sinn et al. 2008) and predators (Grandinson 2005). However, a mothers’ ability to exhibit maternal care and subsequent offspring outcomes, may be mediated by her social relationships. In free-ranging baboons *(Papio cynocephalus ursine*) the offspring of mothers who formed stronger social bonds with maternally related females lived significantly longer, independent of maternal dominance rank (Silk et al. 2009). Female social integration also reduces male harassment in feral horses, leading to increased foal birth rates and survival (Cameron et al. 2009).

When living in groups, maternal care may also be supplemented with alloparental care, in which a non-parent group member provides care for non-descendent young (Riedman 1982). In its most extreme, alloparental care takes the form of cooperative breeding, where nonbreeding helpers raise the young produced by dominant individuals who maintain breeding exclusivity (Jarvis 1981, Lennartz et al. 1987, Clutton-Brock 2002, da Silva Mota et al. 2006). This system dramatically increases juvenile survival (Blumstein and Armitage 1999). In cichlids (*Neolamprologus pulcher*), a cooperative breeding fish, breeding individuals are more connected within their social network than helpers (Dey et al. 2013), indicating that a breeding female’s relationships may work to enforce breeding exclusivity by allowing the dominant individual to police the behaviors of helpers, ensuring better offspring care and outcomes. Clearly there exists a complex relationship between maternal social relationships, maternal care, and offspring outcomes in a variety of species.

Yellow-bellied marmots (*Marmota flaviventer*) (henceforth, ‘marmots’), a large, ground-dwelling alpine rodent (Armitage 2014) classified as semi-cooperative breeders (Blumstein and Armitage 1999), are an ideal species within which to study this phenomena. Marmots are considered facultatively social animals (Blumstein and Armitage 1999) due to the phenotypic plasticity of their sociality; they are capable of both solitary and group living (Huang et al. 2011). Marmot groups vary in size, age, sex demographics and social dynamics (le Roux et al. 2008), and vary in their degree of social interaction according to environmental conditions (Barash 1973). Facultatively social individuals do not need to be social to survive, and therefore have chosen to interact with others either due to some beneficial aspect of their relationships, or because they were forced into group living due to environmental constraints.

Previous work found that more social yearling and adult females who live in large groups were more likely to survive the summer, showing that an individual’s security is affected by their relationships (Montero et al. 2019). Marmots also partially fulfill Solomon and French’s (1997) three cooperative breeding criteria: 1) Pups postpone dispersal past reproductive maturity to the yearling stage; 2) Individuals provide alloparental care in the form of alarm calling, group hibernation, and communal nursing; and 3) There is some degree of reproductive suppression of subordinate females, although more than a single matrilineal female breeds (Blumstein and Armitage 1999). However, none of this breeding behavior is obligate and marmots can live and successfully reproduce in groups that fit none of these criteria (Blumstein and Armitage 1999). This gives us the opportunity to study marmot social groups which exhibit varying degrees of cooperative breeding and draw conclusions about the benefits that maternal social relationships confer on offspring.

Despite the social nature of their reproductive system, marmots seem to experience severe costs of sociality. More social marmots have decreased longevity (Blumstein et al. 2018) and, in particular, increased over-winter mortality (Yang et al. 2017). Additionally, more social females have reduced reproductive success (Wey and Blumstein 2010), a characteristic of reproductive suppression. Adult females (many of whom are mothers) continue to engage in potentially costly affiliative relationships even when they themselves only benefit from group size (Montero et al. 2019). This suggests that other group members (perhaps their offspring) may receive the benefits of sociality. Additionally, while the interaction of sociality and survival has been analyzed in yearlings and adults, it has not been addressed for pups.

Thus, to determine whether maternal relationships benefit pups, we focus here on how maternal social network position is associated with offspring survival. We expect that pups born to more affiliatively socially integrated mothers have a higher likelihood of both summer and yearly survival. Conversely, we expect that pups born to mothers who are more integrated in agonistic networks are less likely to survive. This study investigates a new potential benefit (and cost) of sociality and the effect of social security (Blumstein et al. 2017, Mady and Blumstein 2017, Fuong and Blumstein 2019) in marmot social groups, with potential implications for other cooperative breeding and facultatively social species.

## Methods

### 1. The study system

We studied marmots in and around the Rocky Mountain Biological Laboratory (RMBL) (38°57’29”N, 106°59’06”W, elevation ~2890 m), in the Upper East River Valley in Gunnison County, Colorado, USA. The research was performed with approvals from the UCLA Animal Care Committee (research protocol ARC 2001-191-01) and permits issued from the Colorado Division of Wildlife (TR-519). The marmot population at RMBL has been studied since 1962 (Armitage 2014), and therefore we have long-term data on individuals’ births and deaths (allowing us to calculate accurate ages) and group and colony lineages. A colony is a geographic location that contains one or more marmot social groups. Social groups consist of resident adult females who are maternally related, yearlings (one year old juveniles), pups (juveniles <4 months old), and one or more adult males (Armitage 1991). Three to 4 week old pups emerge from the natal burrow in July (Armitage 1987). Litters average approximately 4 pups at emergence but can range from 1 to 9 pups. Pup survival differs between colonies, but on average roughly 50% of juveniles survive to age 1 year (Schwartz et al. 1998).

Our analysis focuses on data collected between 2002 and 2017. We focused on four colony sites: Bench-River, the Gothic Townsite, Picnic, and Marmot Meadow. Bench-River and the Gothic Townsite are lower elevation sites and we refer to them as “down-valley.” Picnic and Marmot Meadow are higher elevation sites and are therefore considered “up-valley.” Prior work has shown that marmot life history traits, such as body mass, vary as a function of valley position (Van Vuren and Armitage 1991). We selected these sites because we have detailed information on the social interactions that occur at these colonies, these social interactions are highly variable, and the colonies themselves are geographically distinct.

### 2. Trapping

To identify individuals and obtain morphological data and samples, marmots at all four colonies were trapped using Tomahawk live traps baited with horse feed. Every individual at each colony was trapped at least once during the active season (mid-April to mid-September) each year (trapping procedures described in more detail in Armitage 1982). The first time an individual was trapped they were given metal ear tags for permanent identification and temporary dorsal pelage dye marks (using nontoxic Nyanzol-D dye), for observational identification. Hair samples were also collected from trapped individuals for later parentage assignment (Blumstein et al. 2010), and reproductive status noted (more detailed trapping procedures outlined in Armitage 1982).

### 3. Behavioral observations

To collect data on marmot social interactions, near daily observations were performed during periods of peak activity (0700-1000 h and 1600-1900 h), weather permitting, during the active season. Researchers sat at distances that did not influence marmot behavior and observed marmots using binoculars and 15-45 x spotting scopes. Individual marmots were identified using dorsal pelage marks and all social behaviors were recorded (ethogram in Blumstein et al. 2009). The ethogram divides interactions into agonistic interactions (fighting and displacements) or affiliative interactions (all other interactions, excluding sex). Researchers also noted the initiator and receiver of each interaction and the presence of predators (coyotes (*Canis latrans)*, badgers (*Taxidea taxus*), bears (*Ursus arctos* and *Ursus americanus*), red foxes (*Vulpes vulpes)*, golden eagles (*Aquila chrysaetos*), red-tailed hawks (*Buteo jamaicensis)*, other raptors, and ravens (*Corvus corax*)) at a colony during each observation period.

### 4. Determining maternity

To assign maternity to pups we used behavioral and genetic methods. Pups were trapped when they first emerged from the natal burrow and following emergence, predominately engaged in affiliative behaviors with their mothers. This, in conjunction with the identification of lactating mothers during trapping reproductive status assessment, allowed us to behaviorally match pups to mothers. DNA extracted from hair samples collected during trapping confirmed maternity (details in Blumstein et al. 2010).

### 5. Quantifying social relationships

To quantify the social relationships of marmot mothers, we constructed affiliative and agonistic interaction matrices for each of the four colonies for each of the sixteen years of study (agonistic interactions: n = 3,902, affiliative interactions: n = 31,630). Using these matrices, we calculated two sets of ten social network measures (one set based on agonistic interactions and the other set based on affiliative interactions) for all mothers for each year.

The measures we calculated were indegree, outdegree, incloseness, outcloseness, betweenness centrality, eigenvector centrality, instrength, outstrength, negative average shortest path length, and local clustering. Indegree, based on received interactions, and outdegree, based on initiated interactions, are calculated based on the number of individuals with whom a focal individual interacts. These measures gage how directly an individual is connected to others in their network (Wasserman and Faust 1994). If maternal connectivity is positively associated with pup survival, the offspring of mothers with higher indegree and outdegree measures will have higher rates of survival. Closeness centrality, measured as incloseness (received interactions) and outcloseness (initiated interactions), describes how influential a focal individual is within the network, and is calculated as the sum of the reciprocal of the shortest path length between the focal individual and all other individuals in the network (Wasserman and Faust 1994). We predict that more influential mothers will have offspring with greater chances of survival. An individual’s centrality can also be measured as betweenness, which is the proportion of shortest path lengths between all pairs of individuals within the network that include the focal individual (Wey et al. 2008). Similar to closeness centrality, if maternal sociality is positively correlated with offspring survival, the offspring of more central mothers will have a greater chance of survival. Eigenvector centrality is another measure of connectedness, and it takes into account the indirect effects of relationships that occur between a focal individual’s neighbors in addition to the focal individual’s own position (Newman 2018). If a mother has a high degree of eigenvector centrality, her connections are highly connected, which we predict will increase her pups’ chances of survival. Strength, measured as instrength (received interactions) and outstrength (initiated interactions), describes how frequently a focal individual interacts with their neighbors (Wasserman and Faust 1994). We hypothesize that stronger maternal relationships will result in greater pup survival. Average shortest path length is calculated as the average number of individuals in a path between the focal individual and another individual. It describes how efficiently a member of a network can connect with others (Newman 2018). If maternal connection is beneficial for pup survival, a more connected individual should experience greater offspring survival. For the purposes of this analysis we will be using negative average shortest path length, which allows a positive relationship between social integration and pup survival to result in a positive estimate. To understand the cliquishness of a network, clustering calculations generally divide the number of actual relationships formed by a focal individual by the total number of possible relationships (Wey et al. 2008). Local clustering describes an individual’s embeddedness within their network (Wasserman and Faust 1994). We predict that local clustering will have a positive effect on pup survival, because more embedded marmots may receive more help from their conspecifics.

### 6. Quantifying pup survival

We calculated pup survival in the active season and during the pup’s first year of life. Determination of active season survival was based on the date of last sighting for pups known to survive the summer (see also Montero et al. 2019). After pooling all sixteen years of data, 75% of pups known to survive the summer were seen for the last time after 10 August. Therefore, we considered any pup seen after 10 August to have survived the summer. Additionally, any pup who was last seen before 10 August but who reappeared in successive years was also considered to have survived the summer. To calculate yearly survival, any pup who was seen again the following spring after emergence, survived their first year.

### 7. Data processing and analyses

To account for key aspects of a marmot’s environment that are not captured by social network measures, we calculated predator index and social network size. Predator index was based on the proportion of observation periods during which a predator was sighted in a given colony. We used a median split to define a binary measure of comparatively high and low predation pressure for each colony annually. Network size was calculated as the number of individuals in each social network.

To determine the effect of a mother’s social network measures on pup survival, we fitted a series of generalized linear mixed effects models (GLMM) with binomial error structures. Due to the different effects of social network measures on individuals, we fitted a separate model for each social network measure in the agonistic and affiliative iterations and for both summer and yearly survival, giving a total of 40 models. We included as fixed effects the focal social network measure, maternal age, valley position, predation index, network size, pup emergence date, litter size, log of August mass (for yearly survival models) or log of daily mass gain (for summer survival), an interaction term between the focal social network measure and valley position, and an interaction term between the focal social network measure and predation index. We included year and maternal identity as random effects.

We fitted the models in R (R version 3.5.2 2018) using the packages MuMin (Bartoń 2019), lme4 (Bates et al. 2017), lmerTest (Kuznetsova et al. 2016), and optimx (Nash and Varadhan 2011). We used the bobyqa optimizer for some models to obtain proper model convergence.

## Results

### 1. Impact of Maternal Affiliative Interactions on Offspring Summer Survival

In the summer survival model based on affiliative interactions, 8 of the 10 models revealed negative associations with affiliative social relationships. Of these, only instrength was significant at the 0.05 level (Estimate = −0.027, SE = 0.013, z = −2.037, *P* = 0.042; Table 1; Figure 1a). Additionally, some models had significant interactions between the affiliative social network measure and predation index or valley position. In the instrength model (Estimate = 0.030, SE = 0.015, z = 1.986, *P* = 0.047), the indegree model (Estimate = 3.183, SE = 1.599, z = 1.991, *P* = 0.046), the outcloseness model (Estimate = 6.837, SE = 2.408, z = 2.839, *P* = 0.005), and the local clustering model (Estimate = 2.480, SE = 1.162, z = 2.134, *P* = 0.033, the social measure-predation index interaction showed a significant negative effect (Figure 1b, c, d, e), such that the above maternal social measures were positively correlated with offspring success in low predation environments. The interactions between valley position and outcloseness (Estimate = −5.099, SE = 2.191, z = −2.328, *P* = 0.020) and valley position and negative average shortest path length (Estimate = −0.887, SE = 0.446, z = −1.990, *P* = 0.047) had a negative association with pup survival, indicating that maternal out closeness and negative shortest path length are negatively correlated with offspring survival in Picnic and Marmot Meadow (up valley) (Figure 1f, g). Up valley position was a significant positive predictor of offspring survival in 9 models. Daily mass gain was positively significant in all 10 models. Predation index was significant in 7 models, where survival probability decreased with higher predation index. Other fixed effects included in the affiliative summer survival models were not significant (see Supplementary Table S1 for full model results).

**Table 1.**
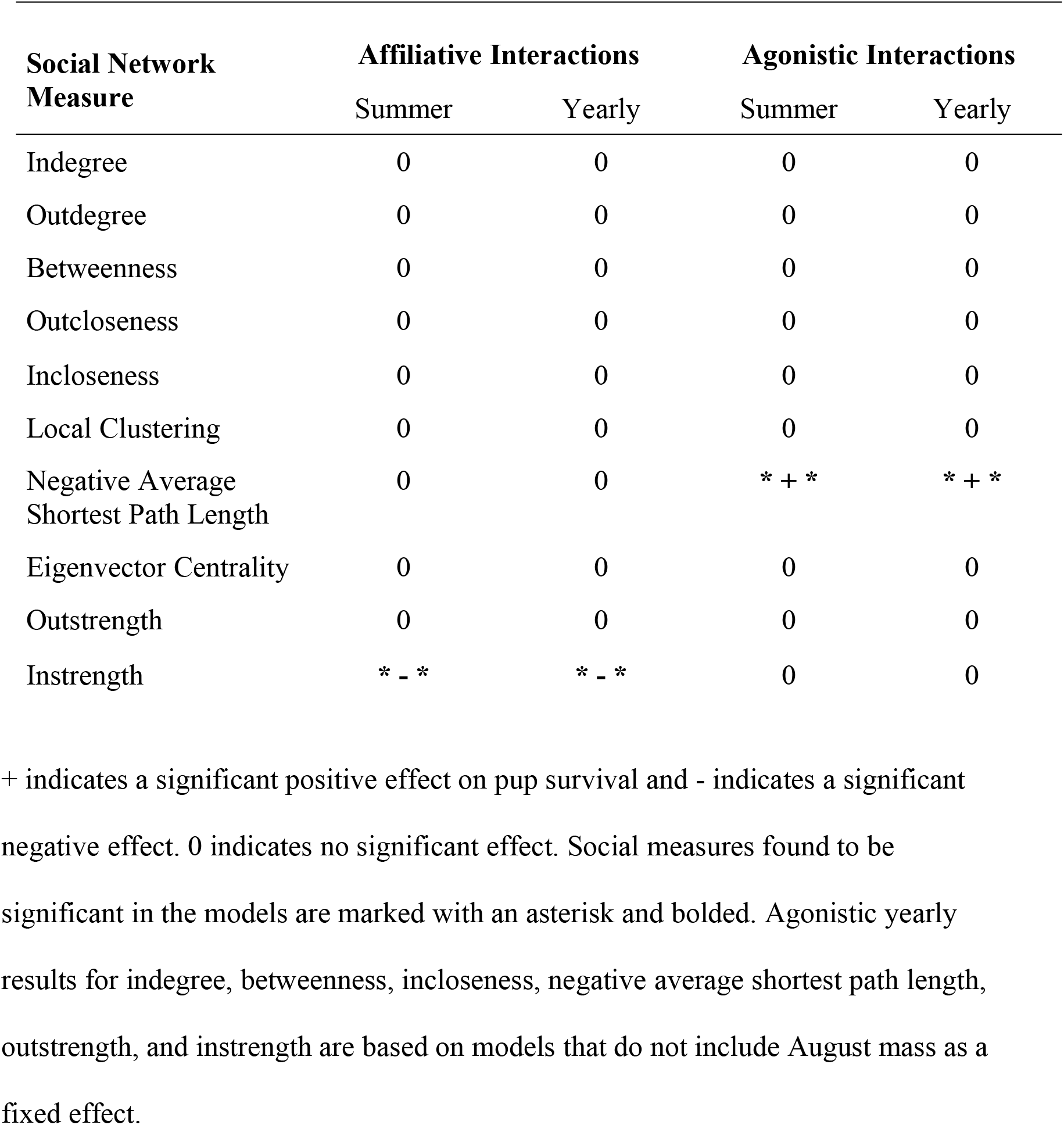
Association between maternal social network measures and pup survival, broken down by summer and yearly survival and affiliative and agonistic networks.

**Figure 1.**
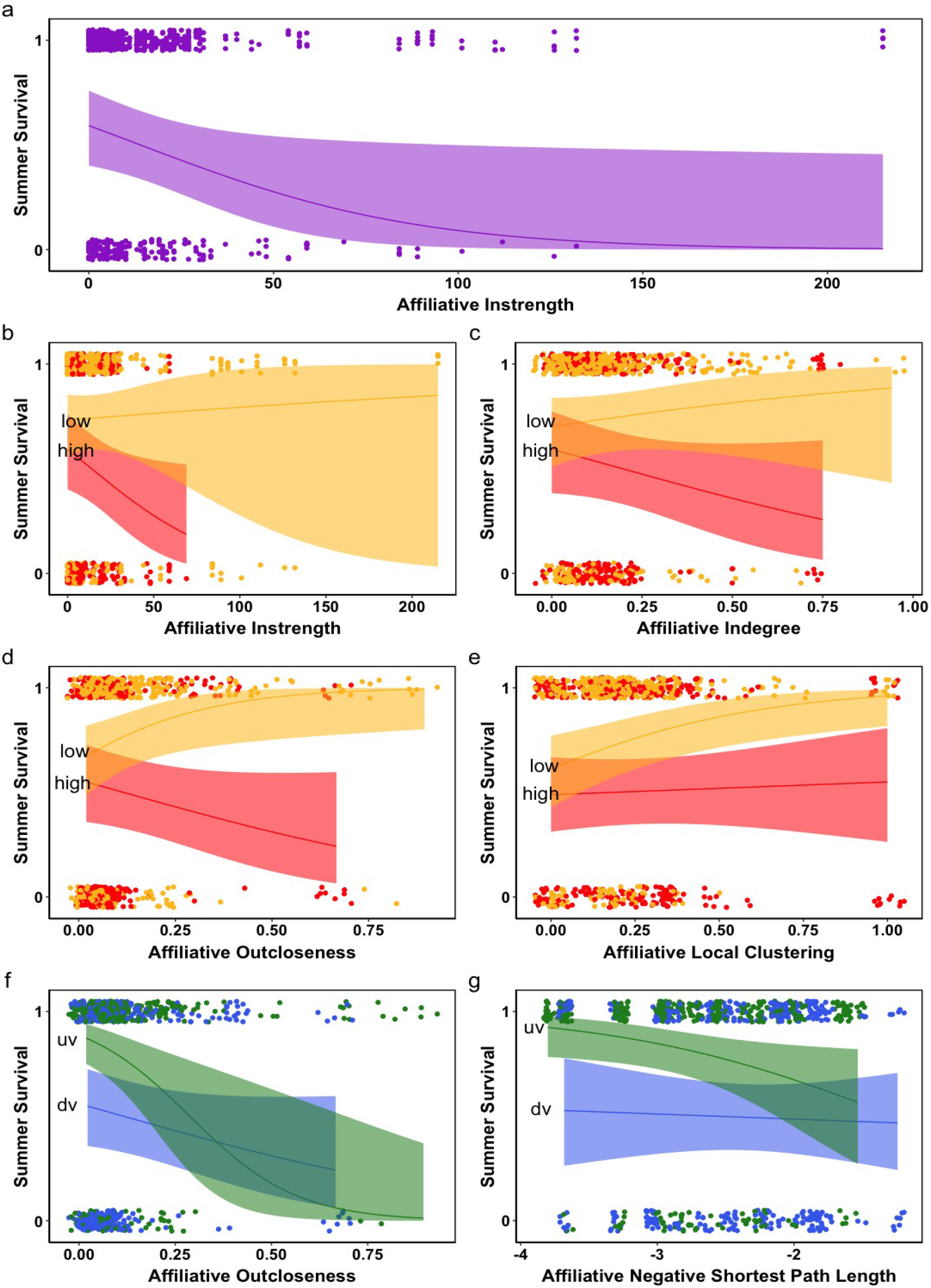
Figures illustrate significant relationships between maternal social network measures and pup summer survival based on affiliative interactions. **a**. Instrength, **b**. Instrength*predation index interaction, **c**. Indegree*predation index interaction, **d**. Outcloseness*predation index interaction, **e**. Local clustering*predation index interaction, **f**. Outcloseness*valley position interaction, **g**. Negative average shortest path length* valley position interaction. Each point represents a pup’s binary survival and the pup’s mother’s social network measure strength. Solid lines show model predictions and shading depicts the 95% confidence interval. Lines in graphs b. – e. depict model predictions for low predation (low) and high predation (high) environments. Lines in graphs f. and g. model predictions for up valley (uv) and down valley (dv) environments. See text for a full description of social measures.

### 2. Impact of Maternal Affiliative Interactions on Offspring Yearly Survival

Six of the 10 yearly survival models based on affiliative interactions had a negative association between affiliative social measure and offspring survival, and 4 of the 10 models were positively associated. None of the positively associated models were significant and, of the models showing a negative association, only instrength was significant (Estimate = −0.031, SE = 0.014, z = −2.263, *P* = 0.024; Table 1; Figure 2a). Up valley position was a significant positive predictor of offspring survival in 9 models. August mass was positively significant in all 10 models. Predation index was significant in 8 models, where survival probability decreased with higher predation index. Both emergence date and network size were positively significant in 1 model. Other fixed effects included in the affiliative yearly survival models were not significant (see Supplementary Table S2 for full model results).

**Figure 2.**
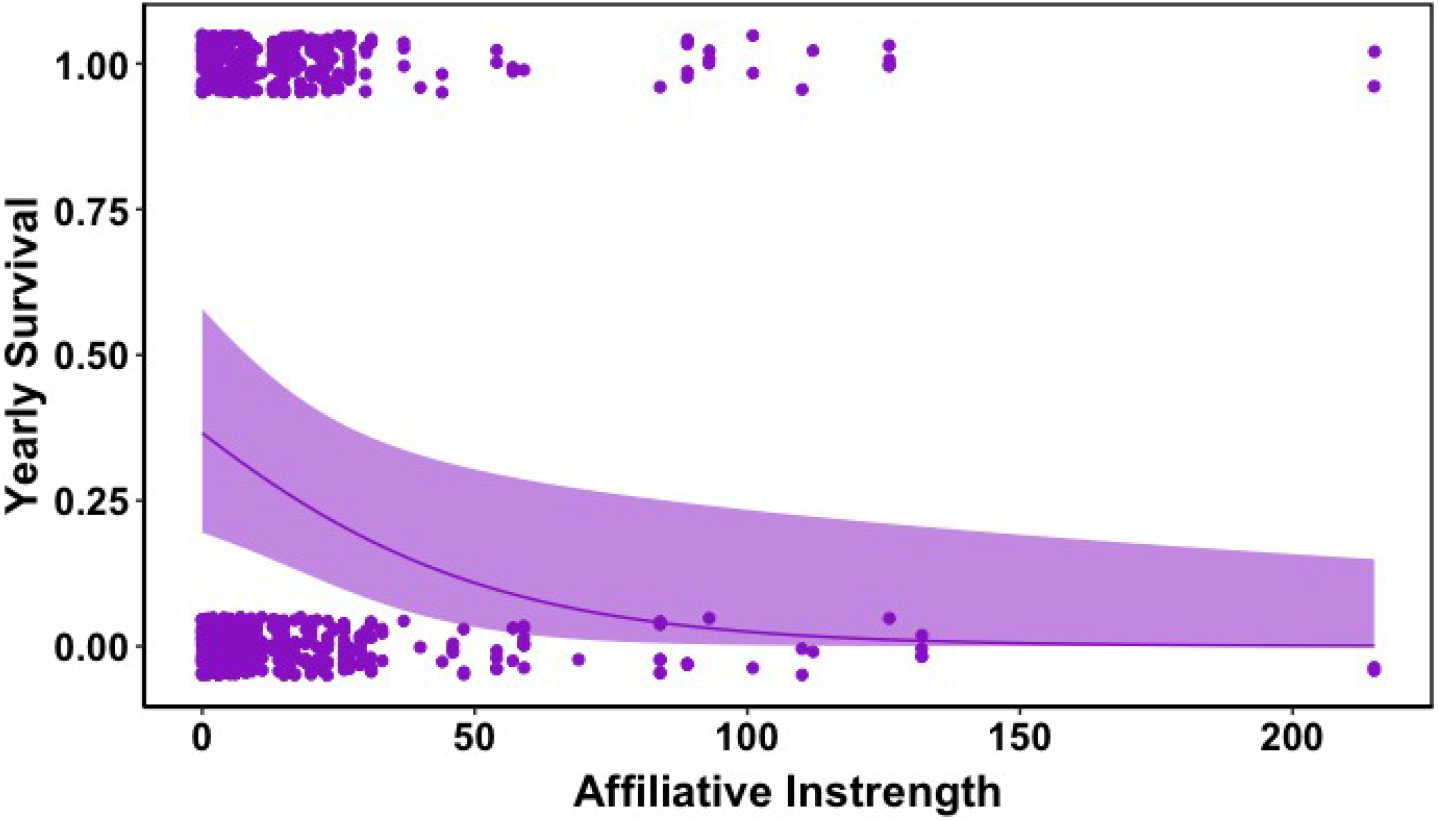
Figure illustrates significant relationships between maternal social network measure instrength and pup yearly survival based on affiliative interactions. Each point in represents a pup’s binary survival and the pup’s mother’s social network measure strength. The solid line shows the model predictions and shading depicts the 95% confidence interval. See text for a full description of social measures.

### 3. Impact of Maternal Agonistic Interactions on Offspring Summer Survival

Four of the 10 summer survival models based on agonistic interactions revealed negative associations with agonistic behavior while the other 6 models had positive associations. Of these models, 1 was significant: negative average shortest path length (Estimate = 1.362, SE = 0.534, z = 2.549, *P* =0.011; Table 1; Figure 3a). In three models, negative average shortest path length (Estimate = −1.917, SE = 0.558, z = −3.434, *P* =0.001), outdegree (Estimate = −5.596, SE = 1.897, z = −2.950, *P* = 0.003), and outcloseness (Estimate = −3.149, SE = 1.552, z = −2.029, *P* = 0.043), there was also a significant negative effect of the interaction of valley position and social network measures. Higher measure of these 3 social measures therefore have a significant negative effect on offspring survival in up valley environments (Figure 3b, c, d). Up valley position was a significant positive predictor of offspring survival in 9 models and a significant negative predictor off offspring survival in 1 model. Daily mass gain was positively significant in 9 models. Predation index was significant in 8 models, where survival probability decreased with higher predation index. Other fixed effects included in the agonistic summer survival models were not significant (see Supplementary Table S3 for full model results).

**Figure 3.**
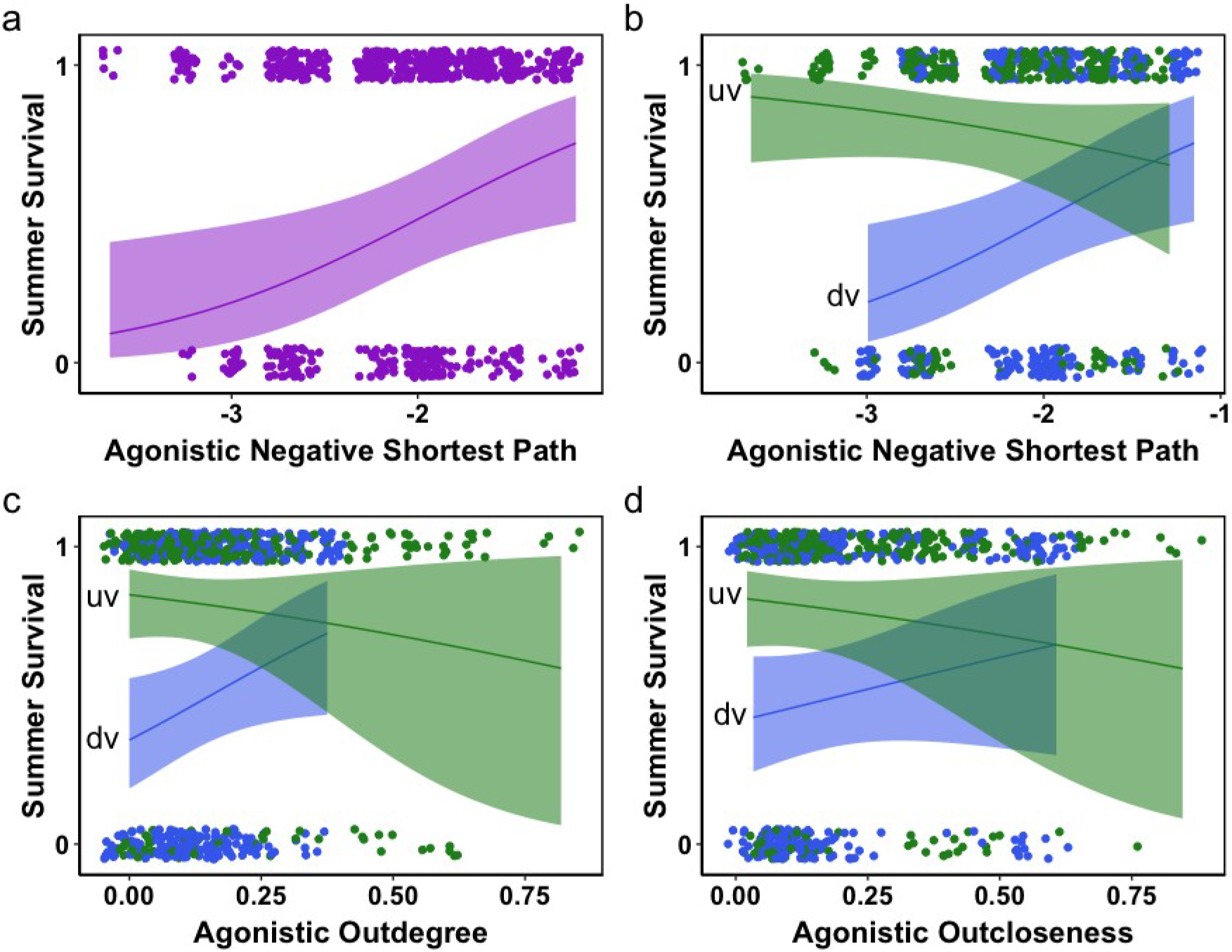
Figures illustrate significant relationships between maternal social network measures and pup summer survival based on agonistic interactions. **a**. Negative average shortest path length, **b**. Negative average shortest path length*valley position interaction, **c**. Outdegree*valley position interaction, **d**. Outcloseness*valley position interaction. Each point represents a pup’s binary survival and the pup’s mother’s social network measure strength. Solid lines show model predictions and shading depicts the 95% confidence interval. Lines in graphs b. – d. depict model predations for up valley (uv) and down valley (dv) environments. See text for a full description of social measures.

### 4. Impact of Maternal Agonistic Interactions on Offspring Yearly Survival

Of the 10 yearly survival models, six models (indegree, betweenness, incloseness, negative average shortest path length, outstrength, and instrength) are singular fitted. Removing august mass eliminates singularity in all six of these models and thus the model results reported for these six social network measures do not include august mass as a fixed effect. Six of the 10 yearly survival models based on agonistic interactions had a positive association between agonistic behavior and pup survival, of which only negative average shortest path length was significant (Estimate = 1.633, SE = 0.537, z = 3.042, *P* = 0.002; Table 1; Figure 4a). There was a significant negative interaction between negative average shortest path length and valley position (Estimate = −1.468, SE = 0.498, z = −2.946, *P* = 0.003), indicating that high measures of maternal negative average shortest path length have a negative effect on offspring survival in up valley environments (Figure 4b). Up valley position was a significant positive predictor of offspring survival in 9 models. August mass was positively significant in 4 models. Predation index was significant in 8 models, where survival probability decreased with higher predation index. Emergence date was positively significant in 4 models and negatively significant in 1 model. Other fixed effects included in the agonistic yearly survival models were not significant (see Supplementary Table S4 for model results).

**Figure 4.**
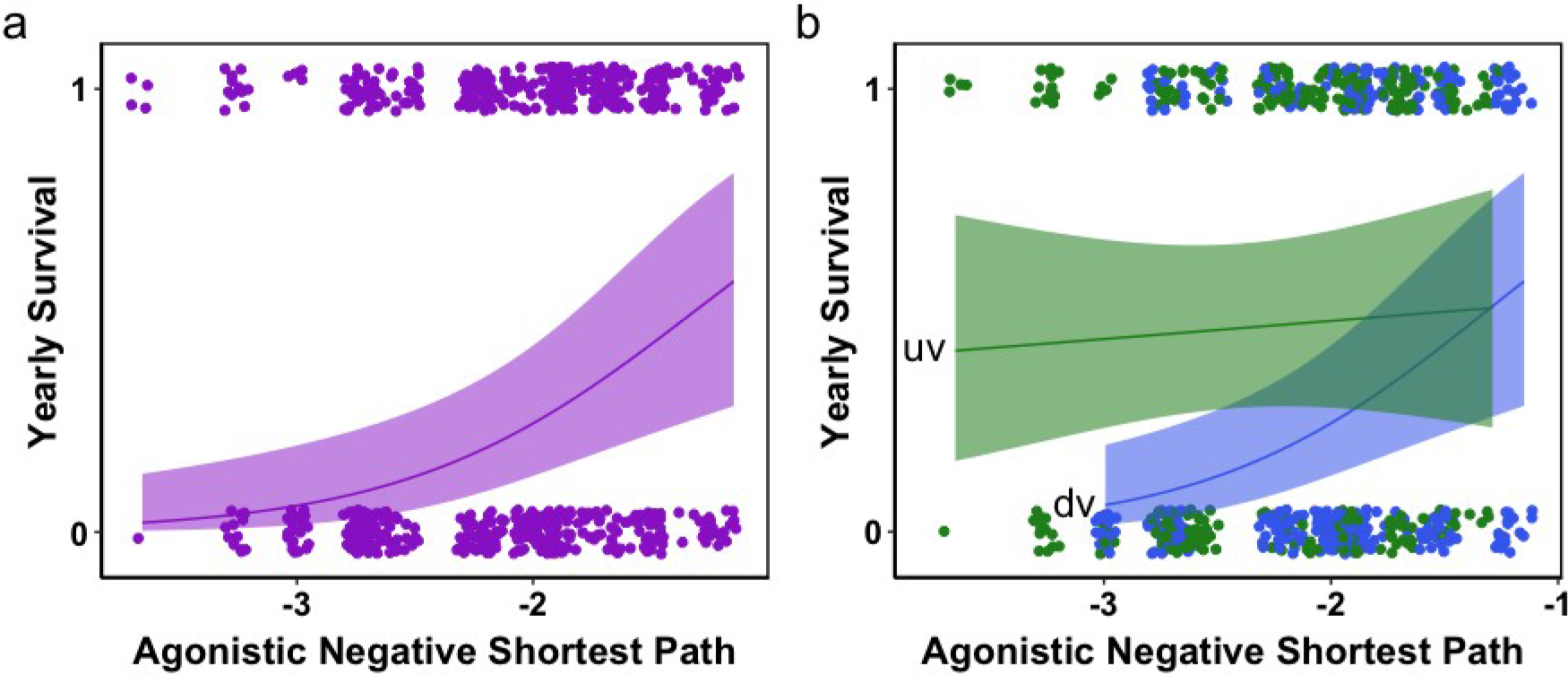
Figures illustrate significant relationships between maternal social network measures and pup yearly survival based on agonistic interactions. **a**. Negative average shortest path length, **b**. Negative average shortest path length*valley position interaction. Each point represents a pup’s binary survival and the pup’s mother’s social network measure strength. Solid lines show model predictions and shading depicts the 95% confidence interval. Lines in graph b. depicts model predations for up valley (uv) and down valley (dv) environments. See text for a full description of social measures.

## Discussion

We investigated the relationship between maternal social network connectivity and integration on offspring survival. We hypothesized that more socially integrated affiliative mothers would have higher rates of offspring survival, indicating a potential benefit of sociality for these facultatively social mammals. We expected to see the opposite effect among more agonistic mothers. We found complex results that suggest that both specific social attributes and environmental factors affect offspring survival in different ways. Social connectedness is neither universally good nor universally poor for marmots but pups are clearly affected by the social world of their mothers.

The strength of received affiliative relationships is associated with *decreased* pup survival over the first year. Conversely, pups are also more likely to survive if their mothers are more central in agonistic networks. Additionally, agonistic negative average shortest path length, rather than the occurrence or frequency of interactions, significantly explains variation in pup survival. This indicates that the pups of mothers who are highly connected to others through agonistic behavior are most successful. At first glance, these findings are contrary to our hypothesis; individuals are not able to seek out better outcomes for their offspring by maintaining strong affiliative relationships that could encourage alloparental care. However, mothers may encourage conspecific help through agonistic behavior, where agonism acts as a punishment intended to enforce altruistic alloparental behavior (in naked mole rats, *Heterocephalus glaber*, Reeve 1992, Clutton-Brock and Parker 1995).

Dominance, while not included in this analysis, plays an important role in shaping how an individual engages with conspecifics and affects offspring survival in many species (Ellis 1995). Dominant individuals may have access to high quality foraging territory (hyenas, *Crocuta crocuta*, Frank et al. 1995 and chimpanzees, *Pan troglodytes*, Pusey 1997), or they may have subordinates help through babysitting and supplemental feeding of offspring (meerkats, *Suricata suricatta*, Russell et al. 2002). In their study of cichlid social networks, Dey et al. (2013) found that dominant reproductive females (of which there is only one) had higher measures of centrality and overall relationship strength. Although marmot dominance is subtle, more dominant females exhibit more agonistic behavior and have greater reproductive success (Blumstein et al. 2016).

Consistent with Dey et al.’s findings, we found that agonistic negative average shortest path length, a measure of maternal centrality, was significantly associated with increased offspring survival. Outstrength, a measure of relationship strength, was also consistently positively associated with offspring survival in three of the four model types, although not significantly so. Therefore, high measures of negative average path length and outstrength may correlate with more dominant individuals, thus enhancing pup survival. While marmots are generalist herbivores and unlikely to dominate food patches, other resources such as hibernacula are limited and may be dominated by females of higher rank, which would improve overwinter survival (Armitage 2014). Additionally, more dominant females may be better able to recruit yearling females as helpers, perhaps through bullying behavior, as discussed above. Yearling females are more likely to disperse when they are less socially connected to their natal colony (Blumstein et al. 2009). Thus, yearling females may seek to minimize the cost and rate of agonistic interactions from more central dominant females by providing alloparental care in a ‘pay to stay’ strategy.

The four complementary sets of analyses illustrate a divergent pattern, whereby affiliative interactions are costly and agonistic interactions are beneficial. These different findings may reflect alternative strategies individuals use to increase their reproductive success (Mendl et al. 1992, Kinahan and Pillay 2008). More dominant mothers may aim to maximize their agonistic centrality, which may win them alloparental helpers, while more subordinate mothers may decrease their affiliative interactions and aim to be more socially isolated, potentially decreasing bullying and competition.

The divergent effects of agonistic and affiliative maternal relationships are consistent across the first summer and the first year of a pup’s life. Affiliative instrength is significantly negatively associated with offspring summer and yearly survival, while agonistic path length has a significant positive effect. This is particularly remarkable given the different common causes of mortality in the summer (predation, Van Vuren 2001) and the winter (unsuccessful hibernation, Schwartz et al. 1998) and the different traits and abilities needed to survive these periods. Our models may reflect a general reality for marmots, that engaging in agonistic behavior is more beneficial than affiliative behavior. However, given that yearly survival is the sum of summer and winter survival, the results for yearly survival are partly driven by the summer effect.

Survival is further complicated by the strong effects of valley position and predation index. The ten significant interaction terms indicate that a maternal social measure that increases pup survival in one environment may have no effect on or even decrease survival in another environment, highlighting the importance of context on social benefits. For example, agonistic maternal path length has a positive effect on pup survival in the summer survival model, but the interaction between path length and valley position in the same model is negative. This means that agonistic maternal path length actually has a negative effect on pup survival in up valley environments. These results highlight the highly context dependent nature of the positive and negative effects of sociality. Thus, general statements about the benefits and costs of maternal social integration will not be applicable to all marmots in all environments. Rather, it is essential that studies on sociality are placed within an environmental context to accurately interpret their effects.

However, there is some consistency in the direction of the interactions. All significant interactions between predation index and maternal social network measure are positive, meaning that in low predation environments, higher maternal social connectivity and integration in both affiliative or agonistic networks has a more consistent, positive effect on offspring outcomes, compared to high predation environments. Pinho et al.’s (2019) work on fecal glucocorticoids and maternal behavior offers a potential mechanism for this phenomenon. More social mothers in lower predation environments have lower stress levels, and pups born to mothers with lower stress levels are more likely to survive. However, mothers in these environments may have lower stress levels precisely because they benefit from conspecific help through alloparental care, as is the case in humans (Balaji et al. 2007). The effect of interactions between valley position and maternal affiliative and agonistic social network measures is also consistent: in up valley environments, increased maternal sociality decreased pup survival. This may be caused by a combination of the negative influence of sociality on over winter survival (Schwartz et al. 1998) and longer winter conditions up valley (Van Vuren and Armitage 1991, Blumstein et al. 2004).

This study adds to a small but growing body of literature documenting the highly environment-dependent nature of sociality. Within the marmot system, social integration, which generally has a positive effect on yearling summer survival, *negatively* impacts yearling summer survival in high predation conditions (Montero et al. 2019). A variety of studies in other systems have investigated the indirect effects of variation in an individuals’ physical environment on their social behavior. Female oribi (*Ourebia ourebi*) have larger group sizes in areas of higher biomass and better food quality, particularly during the dry season (Brashares and Arcese 2002). Female guanacos (*Lama guanicoe*) show increased rates of aggression in scrubland relative to grassland, regardless of predation risk (Marino 2010). Juvenile chum salmon (*Oncorhynchus keta*) increase schooling when food is spatially dispersed, and decrease agonistic interactions in the presence of predators (Ryer and Olla 1996). These approaches offer insights into the relationship between environmental context and the costs and benefits of sociality. However, in focusing only on behavioral plasticity and neglecting fitness effects, these studies fail to capture the full picture: behavioral change in response to environmental variables may not always be adaptive (Badyaev 2005, Ghalambor et al. 2007) or even be possible (DeWitt et al. 1998). Furthermore, unlike our approach, these studies used relatively crude descriptors of social dynamics, such as group size, cohesion, and agonistic behavior, that do not capture the complexity of an individual’s social relationships or the structure of the overall network (Wey et al. 2008, Kurvers et al. 2014).

We therefore suggest that sociality must be studied in a context-dependent manner. The social network approach, used in this study (whereby individuals’ specific social network measures – calculated using observed social interactions – are analyzed as possible predictors of life history traits) is a promising method to quantify and study an individuals’ social environment (Blumstein 2013). Although using social networks to study context-dependent sociality has largely been applied to marmots (e.g., Wey and Blumstein 2010, Yang et al. 2017, Mady and Blumstein 2017, Blumstein et al. 2018), its use should not be restricted to facultatively social species. Obligately social species likely experience many of the same context dependent sociality costs and benefits as facultative social species (Lacey 2004, Maestripieri and Georgiev 2016), although the costs may be more rare and ultimately outweighed by the benefits (Lutermann et al. 2013). This type of analysis may be particularly fruitful in species where populations experience different predation pressures or have significant life history differences. Moving forward, research on how social animals are affected by their environments will be particularly important as habitats change at unprecedented rates due to human use, fragmentation, and climate change.

## Acknowledgements

We are grateful to all marmoteers who collected data over many years, and the Rocky Mountain Biological Laboratory (RMBL) which has helped facilitate this research.

## Data availability

Code and data is available on OSF https://doi.org/10.17605/OSF.IO/WC3NQ.

## Ethics

All procedures were approved under research protocol ARC 2001-191-01 by the University of California Los Angeles Animal Care Committee on 13 May 2002, and renewed annually, as well as annual permits issued by the Colorado Division of Wildlife (TR-519).

## Author contributions

A.P.M., D.M.W., and D.T.B. conceived the project. All authors collected data and designed the data analyses. A.P.M. analyzed the data and wrote the first draft of the manuscript. All authors contributed to subsequent revisions.

