## Supplementary tables for "The effects of maternal social connectivity and integration on offspring survival in a marmot"

Supplementary tables providing estimates and associated statistics for all models fitted in the manuscript.

**Supplementary Table 1. Affiliative Summer Survival Models**
**Supplementary Table 2. Affiliative yearly Survival Models**
**Supplementary Table 3. Agonistic Summer Survival Models**
**Supplementary Table 4. Agonistic yearly Survival Models**

**Supplementary Table 1. Affiliative Summer Survival Models**

Tables show the complete GLMM fixed and random effects outputs for the summer survival models based on affiliative social measures. Social measures include **a**. indegree, **b.** outdegree, **c.** betweenness, **d.** outcloseness, **e.** incloseness, **f.** local clustering, **g.** negative average shortest path length, **h.** eigenvector centrality, **i.** outstrength, and **j.** instrength. Significant variables are bolded and marked with an asterisk.

a. Indegree Fixed Effects

| | Estimate | Std. Error | z value | Pr(>\|z\|) |
|---|---|---|---|---|
| (Intercept) | -2.079 | 1.170 | -1.776 | 0.076 |
| Indegree | -1.911 | 1.317 | -1.451 | 0.147 |
| **Valley_Position_UpValley** | **1.467** | **0.383** | **3.825** | **1.31e-04 *** |
| Predator_Index_Low | 0.472 | 0.353 | 1.336 | 0.182 |
| Pup_Sex_M | 0.062 | 0.203 | 0.308 | 0.758 |
| Litter_Size | -0.036 | 0.060 | -0.608 | 0.543 |
| Network_Size | 0.006 | 0.014 | 0.473 | 0.636 |
| Mother_Age | 0.001 | 0.046 | 0.024 | 0.980 |
| Emergence_Date | 0.016 | 0.017 | 0.951 | 0.341 |
| **Log_Daily_Mass_Gain** | **0.390** | **0.135** | **2.878** | **0.004 *** |
| **Indegree*Predator_Index_Low** | **3.183** | **1.599** | **1.991** | **0.046 *** |
| Indegree* Valley_Position_UpValley | -0.557 | 1.561 | -0.357 | 0.721 |

Indegree Random Effects

| Groups Name | Variance | Std. Dev. |
|---|---|---|
| Mother_UID | 0.139 | 0.372 |
| Year | 1.075 | 1.037 |

b. Outdegree Fixed Effects

| | Estimate | Std. Error | z value | Pr(>\|z\|) |
|---|---|---|---|---|
| (Intercept) | -2.092 | 1.143 | -1.831 | 0.067 |
| Outdegree | -1.202 | 1.435 | -0.838 | 0.402 |
| **Valley_Position_UpValley** | **1.717** | **0.362** | **4.742** | **2.12e-06 *** |
| **Predator_Index_Low** | **0.722** | **0.341** | **2.119** | **0.034 *** |
| Pup_Sex_M | 0.040 | 0.201 | 0.197 | 0.844 |
| Litter_Size | -0.055 | 0.058 | -0.937 | 0.348 |
| Network_Size | 0.005 | 0.013 | 0.362 | 0.718 |
| Mother_Age | -0.008 | 0.045 | -0.181 | 0.856 |
| Emergence_Date | 0.011 | 0.016 | 0.668 | 0.504 |
| **Log_Daily_Mass_Gain** | **0.402** | **0.133** | **3.018** | **0.003 *** |
| Outdegree*Predator_Index_Low | 2.438 | 2.019 | 1.208 | 0.227 |
| Outdegree* Valley_Position_UpValley | -2.121 | 1.957 | -1.084 | 0.278 |

Outdegree Random Effects

| Groups Name | Variance | Std. Dev. |
|---|---|---|
| Mother_UID | 0.088 | 0.297 |
| Year | 0.926 | 0.962 |

c. Betweenness Fixed Effects

| | Estimate | Std. Error | z value | Pr(>\|z\|) |
|---|---|---|---|---|
| (Intercept) | -2.130 | 1.114 | -1.912 | 0.056 |
| Betweenness | -1.739 | 2.089 | -0.833 | 0.405 |
| **Valley_Position_UpValley** | **1.373** | **0.308** | **4.450** | **8.60e-06 *** |
| **Predator_Index_Low** | **0.881** | **0.285** | **3.090** | **0.002 *** |
| Pup_Sex_M | 0.065 | 0.200 | 0.323 | 0.747 |
| Litter_Size | -0.034 | 0.059 | -0.571 | 0.568 |
| Network_Size | 0.007 | 0.012 | 0.549 | 0.583 |
| Mother_Age | 2.64e-04 | 0.045 | 0.006 | 0.995 |
| Emergence_Date | 0.010 | 0.017 | 0.615 | 0.538 |
| **Log_Daily_Mass_Gain** | **0.368** | **0.132** | **2.777** | **0.005 *** |
| Betweenness*Predator_Index_Low | 1.348 | 2.265 | 0.595 | 0.552 |
| Betweenness *Valley_Position_UpValley | 1.589 | 2.262 | 0.702 | 0.483 |

Betweenness Random Effects

| Groups Name | Variance | Std. Dev. |
|---|---|---|
| Mother_UID | 0.115 | 0.340 |
| Year | 0.862 | 0.928 |

d. Outcloseness Fixed Effects

| | Estimate | Std. Error | z value | Pr(>\|z\|) |
|---|---|---|---|---|
| (Intercept) | -2.259 | 1.163 | -1.943 | 0.052 |
| Outcloseness | -2.069 | 1.304 | -1.587 | 0.113 |
| **Valley_Position_UpValley** | **1.822** | **0.351** | **5.194** | **2.06e-07 *** |
| Predator_Index_Low | 0.393 | 0.333 | 1.182 | 0.237 |
| Pup_Sex_M | 0.068 | 0.203 | 0.334 | 0.738 |
| Litter_Size | -0.061 | 0.059 | -1.030 | 0.303 |
| Network_Size | 0.006 | 0.013 | 0.446 | 0.656 |
| Mother_Age | -0.009 | 0.045 | -0.202 | 0.839 |
| Emergence_Date | 0.022 | 0.017 | 1.281 | 0.200 |
| **Log_Daily_Mass_Gain** | **0.413** | **0.135** | **3.069** | **0.002 *** |
| **Outcloseness*Predator_Index_Low** | **6.837** | **2.408** | **2.839** | **0.005 *** |
| **Outcloseness *Valley_Position_UpValley** | **-5.099** | **2.191** | **-2.328** | **0.020 *** |

Outcloseness Random Effects

| Groups Name | Variance | Std. Dev. |
|---|---|---|
| Mother_UID | 0.076 | 0.276 |
| Year | 1.198 | 1.095 |

e. Incloseness Fixed Effects

| | Estimate | Std. Error | z value | Pr(>\|z\|) |
|---|---|---|---|---|
| (Intercept) | -2.157 | 1.140 | -1.892 | 0.059 |
| Incloseness | -0.378 | 1.620 | -0.233 | 0.815 |
| **Valley_Position_UpValley** | **1.705** | **0.356** | **4.786** | **1.70e-06 *** |
| **Predator_Index_Low** | **0.901** | **0.342** | **2.635** | **0.008 *** |
| Pup_Sex_M | 0.048 | 0.201 | 0.238 | 0.812 |
| Litter_Size | -0.042 | 0.058 | -0.736 | 0.462 |
| Network_Size | 0.003 | 0.013 | 0.230 | 0.818 |
| Mother_Age | -0.004 | 0.045 | -0.095 | 0.924 |
| Emergence_Date | 0.009 | 0.017 | 0.541 | 0.589 |
| **Log_Daily_Mass_Gain** | **0.388** | **0.132** | **2.934** | **0.003 *** |
| Incloseness*Predator_Index_Low | 1.030 | 1.863 | 0.553 | 0.580 |
| Incloseness *Valley_Position_UpValley | -1.526 | 1.587 | -0.961 | 0.336 |

Incloseness Random Effects

| Groups Name | Variance | Std. Dev. |
|---|---|---|
| Mother_UID | 0.096 | 0.311 |
| Year | 0.855 | 0.925 |

f. Local Clustering Fixed Effects

| | Estimate | Std. Error | z value | Pr(>\|z\|) |
|---|---|---|---|---|
| **(Intercept)** | **-2.633** | **1.171** | **-2.249** | **0.024 *** |
| Local_Clustering | 0.245 | 0.725 | 0.338 | 0.735 |
| **Valley_Position_UpValley** | **1.161** | **0.350** | **3.320** | **0.001 *** |
| Predator_Index_Low | 0.506 | 0.342 | 1.479 | 0.139 |
| Pup_Sex_M | 0.052 | 0.207 | 0.253 | 0.800 |
| Litter_Size | -0.011 | 0.057 | -0.192 | 0.848 |
| Network_Size | 0.017 | 0.013 | 1.281 | 0.200 |
| Mother_Age | 0.006 | 0.045 | 0.125 | 0.900 |
| Emergence_Date | 0.014 | 0.017 | 0.854 | 0.393 |
| **Log_Daily_Mass_Gain** | **0.355** | **0.135** | **2.633** | **0.008 *** |
| **Local_Clustering* Predator_Index_Low** | **2.480** | **1.162** | **2.134** | **0.033 *** |
| Local_Clustering *Valley_Position_UpValley | 1.377 | 1.092 | 1.261 | 0.207 |

Local Clustering Random Effects

| Groups Name | Variance | Std. Dev. |
|---|---|---|
| Mother_UID | 0.033 | 0.181 |
| Year | 0.879 | 0.937 |

g. Negative Average Shortest Path Length Fixed Effects

| | Estimate | Std. Error | z value | Pr(>\|z\|) |
|---|---|---|---|---|
| (Intercept) | -2.063 | 1.356 | -1.522 | 0.128 |
| Neg_Average_Shortest_Path_Length | -0.097 | 0.356 | -0.273 | 0.785 |
| Valley_Position_UpValley | -0.991 | 1.188 | -0.835 | 0.404 |
| **Predator_Index_Low** | **2.318** | **1.090** | **2.125** | **0.034 *** |
| Pup_Sex_M | 0.082 | 0.201 | 0.405 | 0.685 |
| Litter_Size | -0.060 | 0.058 | -1.032 | 0.302 |
| Network_Size | -0.003 | 0.015 | -0.185 | 0.853 |
| Mother_Age | -0.004 | 0.045 | -0.092 | 0.927 |
| Emergence_Date | 0.011 | 0.016 | 0.680 | 0.496 |
| **Log_Daily_Mass_Gain** | **0.356** | **0.133** | **2.680** | **0.007 *** |
| Neg_Average_Shortest_Path_Length *Predator_Index_Low | 0.477 | 0.419 | 1.137 | 0.255 |
| **Neg_Average_Shortest_Path_Length * Valley_Position_UpValley** | **-0.887** | **0.446** | **-1.990** | **0.047 *** |

Negative Average Shortest Path Length Random Effects

| Groups Name | Variance | Std. Dev. |
|---|---|---|
| Mother_UID | 0.094 | 0.307 |
| Year | 0.878 | 0.937 |

h. Eigenvector Centrality Fixed Effects

| | Estimate | Std. Error | z value | Pr(>\|z\|) |
|---|---|---|---|---|
| **(Intercept)** | **-2.335** | **1.1451** | **-2.039** | **0.041 *** |
| Eigenvector_Centrality | -0.007 | 0.547 | -0.013 | 0.989 |
| **Valley_Position_UpValley** | **1.534** | **0.365** | **4.207** | **2.59e-05 *** |
| **Predator_Index_Low** | **0.774** | **0.323** | **2.397** | **0.017 *** |
| Pup_Sex_M | 0.061 | 0.201 | 0.305 | 0.761 |
| Litter_Size | -0.037 | 0.059 | -0.628 | 0.530 |
| Network_Size | 0.009 | 0.013 | 0.724 | 0.469 |
| Mother_Age | -0.007 | 0.046 | -0.145 | 0.885 |
| Emergence_Date | 0.014 | 0.017 | 0.799 | 0.424 |
| **Log_Daily_Mass_Gain** | **0.380** | **0.133** | **2.848** | **0.004 *** |
| Eigenvector_Centrality *Predator_Index_Low | 0.612 | 0.699 | 0.875 | 0.381 |
| Eigenvector_Centrality *Valley_Position_UpValley | -0.296 | 0.682 | -0.434 | 0.664 |

Eigenvector Centrality Random Effects

| Groups Name | Variance | Std. Dev. |
|---|---|---|
| Mother_UID | 0.137 | 0.370 |
| Year | 0.901 | 0.949 |

i. Outstrength Fixed Effects

| | Estimate | Std. Error | z value | Pr(>\|z\|) |
|---|---|---|---|---|
| **(Intercept)** | **-2.407** | **1.101** | **-2.185** | **0.029 *** |
| Outstrength | 0.012 | 0.025 | 0.486 | 0.627 |
| **Valley_Position_UpValley** | **1.893** | **0.313** | **6.038** | **1.56e-09 *** |
| **Predator_Index_Low** | **0.961** | **0.306** | **3.146** | **0.002 *** |
| Pup_Sex_M | 0.036 | 0.199 | 0.180 | 0.857 |
| Litter_Size | -0.050 | 0.056 | -0.887 | 0.375 |
| Network_Size | 0.005 | 0.012 | 0.388 | 0.698 |
| Mother_Age | -0.007 | 0.044 | -0.171 | 0.864 |
| Emergence_Date | 0.007 | 0.016 | 0.447 | 0.655 |
| **Log_Daily_Mass_Gain** | **0.416** | **0.130** | **3.191** | **0.001 *** |
| Outstrength*Predator_Index_Low | 0.010 | 0.025 | 0.390 | 0.697 |
| Outstrength *Valley_Position_UpValley | -0.037 | 0.021 | -1.819 | 0.069 |

Outstrength Random Effects

| Groups Name | Variance | Std. Dev. |
|---|---|---|
| Mother_UID | 0.036 | 0.191 |
| Year | 0.736 | 0.858 |

j. Instrength Fixed Effects

| | Estimate | Std. Error | z value | Pr(>\|z\|) |
|---|---|---|---|---|
| (Intercept) | -1.908 | 1.177 | -1.622 | 0.105 |
| **Instrength** | **-0.027** | **0.013** | **-2.037** | **0.042** * |
| **Valley_Position_UpValley** | **1.715** | **0.334** | **5.126** | **2.95e-07** * |
| **Predator_Index_Low** | **0.633** | **0.321** | **1.971** | **0.049** * |
| Pup_Sex_M | 0.036 | 0.202 | 0.176 | 0.861 |
| Litter_Size | -0.064 | 0.059 | -1.075 | 0.282 |
| Network_Size | 0.007 | 0.012 | 0.604 | 0.546 |
| Mother_Age | -0.004 | 0.045 | -0.085 | 0.933 |
| Emergence_Date | 0.008 | 0.016 | 0.487 | 0.626 |
| **Log_Daily_Mass_Gain** | **0.413** | **0.137** | **3.020** | **0.002** * |
| **Instrength*Predator_Index_Low** | **0.030** | **0.015** | **1.986** | **0.047** * |
| Instrength *Valley_Position_UpValley | -0.012 | 0.013 | -0.886 | 0.376 |

Instrength Random Effects

| Groups Name | Variance | Std. Dev. |
|---|---|---|
| Mother_UID | 0.084 | 0.290 |
| Year | 0.971 | 0.986 |

**Supplementary Table 2. Affiliative Yearly Survival Models**

Tables show the complete GLMM fixed and random effects outputs for the yearly survival models based on affiliative social measures. Social measures include **a**. indegree, **b.** outdegree, **c.** betweenness, **d.** outcloseness, **e.** incloseness, **f.** local clustering, **g.** negative average shortest path length, **h.** eigenvector centrality, **i.** outstrength, and **j.** instrength. Significant variables are bolded and marked with an asterisk.

a. Indegree Fixed Effects

| | Estimate | Std. Error | z value | Pr(>\|z\|) |
|---|---|---|---|---|
| **(Intercept)** | **-7.673** | **1.492** | **-5.142** | **2.72e-07 *** |
| Indegree | -0.776 | 1.355 | -0.573 | 0.567 |
| **Valley_Position_UpValley** | **0.851** | **0.344** | **2.471** | **0.014 *** |
| **Predator_Index_Low** | **0.778** | **0.326** | **2.386** | **0.017 *** |
| Pup_Sex_M | -0.188 | 0.188 | -1.004 | 0.315 |
| Litter_Size | -0.028 | 0.058 | -0.483 | 0.629 |
| Network_Size | 0.025 | 0.013 | 1.910 | 0.056 |
| Mother_Age | -0.003 | 0.044 | -0.062 | 0.951 |
| Emergence_Date | 0.035 | 0.019 | 1.814 | 0.070 |
| **Log_August_Mass** | **1.002** | **0.175** | **5.709** | **1.14e-08 *** |
| Indegree*Predator_Index_Low | 1.537 | 1.525 | 1.008 | 0.314 |
| Indegree* Valley_Position_UpValley | 0.901 | 1.422 | 0.633 | 0.527 |

Indegree Random Effects

| Groups Name | Variance | Std. Dev. |
|---|---|---|
| Mother_UID | 0.159 | 0.399 |
| Year | 1.667 | 1.291 |

b. Outdegree Fixed Effects

| | Estimate | Std. Error | z value | Pr(>\|z\|) |
|---|---|---|---|---|
| **(Intercept)** | **-7.904** | **1.497** | **-5.281** | **1.29e-07 *** |
| Outdegree | 0.571 | 1.577 | 0.362 | 0.717 |
| **Valley_Position_UpValley** | **1.347** | **0.346** | **3.889** | **1.01e-04 *** |
| **Predator_Index_Low** | **0.862** | **0.329** | **2.624** | **0.009 *** |
| Pup_Sex_M | -0.200 | 0.187 | -1.069 | 0.285 |
| Litter_Size | -0.037 | 0.058 | -0.650 | 0.516 |
| Network_Size | 0.022 | 0.013 | 1.758 | 0.079 |
| Mother_Age | -0.008 | 0.045 | -0.183 | 0.854 |
| Emergence_Date | 0.035 | 0.019 | 1.828 | 0.068 |
| **Log_August_Mass** | **1.020** | **0.176** | **5.809** | **6.28e-09 *** |
| Outdegree*Predator_Index_Low | 2.242 | 2.000 | 1.121 | 0.262 |
| Outdegree * Valley_Position_UpValley | -2.619 | 1.916 | -1.367 | 0.172 |

Outdegree Random Effects

| Groups Name | Variance | Std. Dev. |
|---|---|---|
| Mother_UID | 0.158 | 0.397 |
| Year | 1.669 | 1.292 |

c. Betweenness Fixed Effects

| | Estimate | Std. Error | z value | Pr(>\|z\|) |
|---|---|---|---|---|
| **(Intercept)** | **-7.5685** | **1.444** | **-5.240** | **1.61e-07 *** |
| Betweenness | -2.447 | 2.372 | -1.031 | 0.302 |
| **Valley_Position_UpValley** | **1.055** | **0.289** | **3.645** | **2.67e-04 *** |
| **Predator_Index_Low** | **0.883** | **0.277** | **3.195** | **0.001 *** |
| Pup_Sex_M | -0.198 | 0.186 | -1.063 | 0.288 |
| Litter_Size | -0.027 | 0.057 | -0.469 | 0.639 |
| Network_Size | 0.020 | 0.012 | 1.630 | 0.103 |
| Mother_Age | 0.001 | 0.044 | 0.012 | 0.991 |
| Emergence_Date | 0.030 | 0.019 | 1.599 | 0.110 |
| **Log_August_Mass** | **1.007** | **0.173** | **5.805** | **6.43e-09 *** |
| Betweenness*Predator_Index_Low | 2.647 | 2.200 | 1.203 | 0.229 |
| Betweenness *Valley_Position_UpValley | 1.153 | 2.328 | 0.495 | 0.620 |

Betweenness Random Effects

| Groups Name | Variance | Std. Dev. |
|---|---|---|
| Mother_UID | 0.114 | 0.337 |
| Year | 1.561 | 1.249 |

d. Outcloseness Fixed Effects

| | Estimate | Std. Error | z value | Pr(>\|z\|) |
|---|---|---|---|---|
| **(Intercept)** | **-8.057** | **1.498** | **-5.379** | **7.49e-08 *** |
| Outcloseness | 1.185 | 1.322 | 0.896 | 0.370 |
| **Valley_Position_UpValley** | **1.257** | **0.334** | **3.763** | **1.68e-04 *** |
| **Predator_Index_Low** | **0.990** | **0.315** | **3.147** | **0.002 *** |
| Pup_Sex_M | -0.192 | 0.188 | -1.022 | 0.307 |
| Litter_Size | -0.030 | 0.058 | -0.512 | 0.609 |
| Network_Size | 0.023 | 0.012 | 1.849 | 0.065 |
| Mother_Age | -0.007 | 0.045 | -0.149 | 0.881 |
| **Emergence_Date** | **0.037** | **0.020** | **1.887** | **0.059 *** |
| **Log_August_Mass** | **1.014** | **0.175** | **5.775** | **7.68e-09 *** |
| Outcloseness*Predator_Index_Low | 1.375 | 2.215 | 0.621 | 0.535 |
| Outcloseness *Valley_Position_UpValley | -2.116 | 2.044 | -1.035 | 0.301 |

Outcloseness Random Effects

| Groups Name | Variance | Std. Dev. |
|---|---|---|
| Mother_UID | 0.171 | 0.413 |
| Year | 1.545 | 1.243 |

e. Incloseness Fixed Effects

| | Estimate | Std. Error | z value | Pr(>\|z\|) |
|---|---|---|---|---|
| **(Intercept)** | **-7.787** | **1.481** | **-5.257** | **1.47e-07 *** |
| Incloseness | -0.259 | 1.770 | -0.147 | 0.883 |
| **Valley_Position_UpValley** | **1.281** | **0.333** | **3.842** | **1.22e-04 *** |
| **Predator_Index_Low** | **0.814** | **0.318** | **2.561** | **0.010 *** |
| Pup_Sex_M | -0.192 | 0.187 | -1.028 | 0.304 |
| Litter_Size | -0.032 | 0.057 | -0.556 | 0.578 |
| Network_Size | 0.020 | 0.012 | 1.620 | 0.105 |
| Mother_Age | -0.005 | 0.044 | -0.115 | 0.909 |
| Emergence_Date | 0.034 | 0.019 | 1.805 | 0.071 |
| **Log_August_Mass** | **1.0160** | **0.173** | **5.862** | **4.58e-09 *** |
| Incloseness*Predator_Index_Low | 2.582 | 1.958 | 1.319 | 0.187 |
| Incloseness *Valley_Position_UpValley | -2.031 | 1.640 | -1.238 | 0.216 |

Incloseness Random Effects

| Groups Name | Variance | Std. Dev. |
|---|---|---|
| Mother_UID | 0.142 | 0.376 |
| Year | 1.620 | 1.273 |

f. Local Clustering Fixed Effects

| | Estimate | Std. Error | z value | Pr(>\|z\|) |
|---|---|---|---|---|
| **(Intercept)** | **-7.441** | **1.517** | **-4.904** | **9.39e-07 *** |
| Local_Clustering | 0.282 | 0.845 | 0.334 | 0.739 |
| **Valley_Position_UpValley** | **1.279** | **0.366** | **3.490** | **4.82e-04 *** |
| Predator_Index_Low | 0.560 | 0.352 | 1.591 | 0.112 |
| Pup_Sex_M | -0.184 | 0.194 | -0.947 | 0.343 |
| Litter_Size | -0.026 | 0.057 | -0.455 | 0.649 |
| **Network_Size** | **0.026** | **0.013** | **2.052** | **0.040 *** |
| Mother_Age | 0.015 | 0.045 | 0.328 | 0.743 |
| Emergence_Date | 0.030 | 0.020 | 1.551 | 0.121 |
| **Log_August_Mass** | **0.918** | **0.178** | **5.148** | **2.63e-07 *** |
| Local_Clustering *Predator_Index_Low | 2.028 | 1.099 | 1.846 | 0.065 |
| Local_Clustering *Valley_Position_UpValley | -0.541 | 1.116 | -0.485 | 0.628 |

Local Clustering Random Effects

| Groups Name | Variance | Std. Dev. |
|---|---|---|
| Mother_UID | 0.133 | 0.364 |
| Year | 1.497 | 1.223 |

g. Negative Average Shortest Path Length Fixed Effects

| | Estimate | Std. Error | z value | Pr(>\|z\|) |
|---|---|---|---|---|
| **(Intercept)** | **-6.249** | **1.701** | **-3.674** | **2.39e-04 *** |
| Neg_Average_Shortest_Path_Length | 0.522 | 0.377 | 1.384 | 0.166 |
| Valley_Position_UpValley | -0.028 | 1.033 | -0.027 | 0.978 |
| Predator_Index_Low | 0.220 | 0.963 | 0.229 | 0.819 |
| Pup_Sex_M | -0.189 | 0.187 | -1.009 | 0.313 |
| Litter_Size | -0.051 | 0.057 | -0.894 | 0.371 |
| Network_Size | 0.021 | 0.015 | 1.380 | 0.167 |
| Mother_Age | 0.001 | 0.044 | 0.033 | 0.974 |
| Emergence_Date | 0.031 | 0.019 | 1.599 | 0.110 |
| **Log_August_Mass** | **0.988** | **0.175** | **5.628** | **1.83e-08 *** |
| Neg_Average_Shortest_Path_Length *Predator_Index_Low | -0.370 | 0.360 | -1.027 | 0.304 |
| Neg_Average_Shortest_Path_Length *Valley_Position_UpValley | -0.466 | 0.384 | -1.216 | 0.224 |

Negative Average Shortest Path Length Random Effects

| Groups Name | Variance | Std. Dev. |
|---|---|---|
| Mother_UID | 0.132 | 0.363 |
| Year | 1.720 | 1.311 |

h. Eigenvector Centrality Fixed Effects

| | Estimate | Std. Error | z value | Pr(>\|z\|) |
|---|---|---|---|---|
| **(Intercept)** | **-7.467** | **1.471** | **-5.077** | **3.84e-07 *** |
| Eigenvector_Centrality | -0.723 | 0.611 | -1.183 | 0.237 |
| **Valley_Position_UpValley** | **1.010** | **0.340** | **2.970** | **0.003 *** |
| **Predator_Index_Low** | **0.751** | **0.307** | **2.448** | **0.014 *** |
| Pup_Sex_M | -0.191 | 0.187 | -1.024 | 0.306 |
| Litter_Size | -0.041 | 0.057 | -0.718 | 0.473 |
| Network_Size | 0.020 | 0.012 | 1.728 | 0.084 |
| Mother_Age | -0.006 | 0.044 | -0.142 | 0.887 |
| Emergence_Date | 0.037 | 0.019 | 1.921 | 0.055 |
| **Log_August_Mass** | **1.005** | **0.174** | **5.777** | **7.62e-09 *** |
| Eigenvector_Centrality *Predator_Index_Low | 1.003 | 0.673 | 1.490 | 0.136 |
| Eigenvector_Centrality *Valley_Position_UpValley | 0.124 | 0.646 | 0.192 | 0.848 |

Eigenvector Centrality Random Effects

| Groups Name | Variance | Std. Dev. |
|---|---|---|
| Mother_UID | 0.132 | 0.364 |
| Year | 1.721 | 1.312 |

i. Outstrength Fixed Effects

| | Estimate | Std. Error | z value | Pr(>\|z\|) |
|---|---|---|---|---|
| **(Intercept)** | **-7.456** | **1.455** | **-5.123** | **3.01e-07 *** |
| Outstrength | -0.016 | 0.025 | -0.626 | 0.531 |
| **Valley_Position_UpValley** | **1.368** | **0.298** | **4.584** | **4.56e-06 *** |
| **Predator_Index_Low** | **0.973** | **0.294** | **3.312** | **0.000925 *** |
| Pup_Sex_M | -0.185 | 0.186 | -0.992 | 0.321 |
| Litter_Size | -0.054 | 0.057 | -0.946 | 0.344 |
| Network_Size | 0.018 | 0.012 | 1.541 | 0.123 |
| Mother_Age | -0.006 | 0.044 | -0.140 | 0.889 |
| Emergence_Date | 0.028 | 0.019 | 1.486 | 0.137 |
| **Log_August_Mass** | **1.020** | **0.173** | **5.903** | **3.57e-09 *** |
| Outstrength*Predator_Index_Low | 0.021 | 0.025 | 0.823 | 0.410 |
| Outstrength *Valley_Position_UpValley | -0.018 | 0.015 | -1.143 | 0.253 |

Outstrength Random Effects

| Groups Name | Variance | Std. Dev. |
|---|---|---|
| Mother_UID | 0.119 | 0.345 |
| Year | 1.495 | 1.223 |

j. Instrength Fixed Effects

<table>
<tr><th></th><th>Estimate</th><th>Std. Error</th><th>z value</th><th>Pr(>|z|)</th></tr>
<tr><td>(Intercept)</td><td>-6.750</td><td>1.438</td><td>-4.694</td><td>2.68e-06 *</td></tr>
<tr><td>Instrength</td><td>-0.031</td><td>0.014</td><td>-2.263</td><td>0.024 *</td></tr>
<tr><td>Valley_Position_UpValley</td><td>1.028</td><td>0.293</td><td>3.504</td><td>4.58e-04 *</td></tr>
<tr><td>Predator_Index_Low</td><td>0.831</td><td>0.285</td><td>2.913</td><td>0.004 *</td></tr>
<tr><td>Pup_Sex_M</td><td>-0.192</td><td>0.185</td><td>-1.039</td><td>0.299</td></tr>
<tr><td>Litter_Size</td><td>-0.070</td><td>0.056</td><td>-1.250</td><td>0.211</td></tr>
<tr><td>Network_Size</td><td>0.017</td><td>0.011</td><td>1.482</td><td>0.138</td></tr>
<tr><td>Mother_Age</td><td>0.003</td><td>0.042</td><td>0.074</td><td>0.941</td></tr>
<tr><td>Emergence_Date</td><td>0.028</td><td>0.018</td><td>1.539</td><td>0.124</td></tr>
<tr><td>Log_August_Mass</td><td>0.978</td><td>0.168</td><td>5.825</td><td>5.70e-09 *</td></tr>
<tr><td>Instrength*Predator_Index_Low</td><td>0.018</td><td>0.015</td><td>1.226</td><td>0.220</td></tr>
<tr><td>Instrength *Valley_Position_UpValley</td><td>0.007</td><td>0.010</td><td>0.729</td><td>0.466</td></tr>
</table>

Instrength Random Effects

| Groups Name | Variance | Std. Dev. |
|---|---|---|
| Mother_UID | 0.053 | 0.231 |
| Year | 1.590 | 1.261 |

**Supplementary Table 3. Agonistic Summer Survival Models**

Tables show the complete GLMM fixed and random effects outputs for the summer survival models based on agonistic social measures. Social measures include **a**. indegree, **b.** outdegree, **c.** betweenness, **d.** outcloseness, **e.** incloseness, **f.** local clustering, **g.** average shortest path length, **h.** eigenvector centrality, **i.** outstrength, and **j.** instrength. Significant variables are bolded and marked with an asterisk. Two asterisks (**) signify a p-value < 0.05 and one asterisk (*) signifies a p-value between 0.05 and 0.10.

a. Indegree Fixed Effects

| | Estimate | Std. Error | z value | Pr(>\|z\|) |
|---|---|---|---|---|
| (Intercept) | -1.894 | 1.283 | -1.476 | 0.140 |
| Indegree | -1.773 | 1.967 | -0.902 | 0.367 |
| **Valley_Position_UpValley** | **1.903** | **0.424** | **4.487** | **7.21e-06 *** |
| Predator_Index_Low | 0.801 | 0.421 | 1.903 | 0.057 |
| Pup_Sex_M | 0.011 | 0.214 | 0.050 | 0.960 |
| Litter_Size | -0.026 | 0.063 | -0.405 | 0.686 |
| Network_Size | 0.002 | 0.014 | 0.167 | 0.868 |
| Mother_Age | 0.001 | 0.049 | 0.027 | 0.979 |
| Emergence_Date | 0.013 | 0.018 | 0.720 | 0.471 |
| **Log_Daily_Mass_Gain** | **0.355** | **0.147** | **2.407** | **0.016 *** |
| Indegree*Predator_Index_Low | 2.556 | 2.108 | 1.212 | 0.225 |
| Indegree* Valley_Position_UpValley | -2.450 | 1.743 | -1.406 | 0.160 |

Indegree Random Effects

| Groups Name | Variance | Std. Dev. |
|---|---|---|
| Mother_UID | 0.150 | 0.388 |
| Year | 0.897 | 0.947 |

b. Outdegree Fixed Effects

| | Estimate | Std. Error | z value | Pr(>\|z\|) |
|---|---|---|---|---|
| **(Intercept)** | **-2.979** | **1.260** | **-2.364** | **0.018 *** |
| Outdegree | 3.999 | 2.074 | 1.928 | 0.054 |
| **Valley_Position_UpValley** | **2.256** | **0.400** | **5.641** | **1.69e-08 *** |
| **Predator_Index_Low** | **1.259** | **0.413** | **3.047** | **0.002 *** |
| Pup_Sex_M | 0.006 | 0.215 | 0.028 | 0.977 |
| Litter_Size | -0.024 | 0.063 | -0.371 | 0.710 |
| Network_Size | 0.006 | 0.014 | 0.442 | 0.658 |
| Mother_Age | -0.008 | 0.049 | -0.158 | 0.875 |
| Emergence_Date | 0.016 | 0.018 | 0.901 | 0.367 |
| **Log_Daily_Mass_Gain** | **0.376** | **0.146** | **2.583** | **0.010 *** |
| Outdegree*Predator_Index_Low | -0.333 | 2.254 | -0.148 | 0.883 |
| **Outdegree *Valley_Position_UpValley** | **-5.596** | **1.897** | **-2.950** | **0.003 *** |

Outdegree Random Effects

| Groups Name | Variance | Std. Dev. |
|---|---|---|
| Mother_UID | 0.171 | 0.413 |
| Year | 0.640 | 0.800 |

c. Betweenness Fixed Effects

| | Estimate | Std. Error | z value | Pr(>\|z\|) |
|---|---|---|---|---|
| (Intercept) | -2.104 | 1.213 | -1.735 | 0.083 |
| Betweenness | -3.144 | 2.190 | -1.435 | 0.151 |
| **Valley_Position_UpValley** | **1.321** | **0.318** | **4.154** | **3.27e-05 *** |
| **Predator_Index_Low** | **0.957** | **0.291** | **3.287** | **0.001 *** |
| Pup_Sex_M | 0.041 | 0.214 | 0.190 | 0.849 |
| Litter_Size | 0.006 | 0.064 | 0.092 | 0.926 |
| Network_Size | 0.009 | 0.013 | 0.647 | 0.518 |
| Mother_Age | -0.006 | 0.049 | -0.122 | 0.903 |
| Emergence_Date | 0.011 | 0.018 | 0.591 | 0.555 |
| **Log_Daily_Mass_Gain** | **0.326** | **0.147** | **2.226** | **0.026 *** |
| Betweenness*Predator_Index_Low | 3.442 | 2.759 | 1.248 | 0.212 |
| Betweenness *Valley_Position_UpValley | 3.459 | 3.186 | 1.086 | 0.278 |

Betweenness Random Effects

| Groups Name | Variance | Std. Dev. |
|---|---|---|
| Mother_UID | 0.228 | 0.477 |
| Year | 0.854 | 0.924 |

d. Outcloseness Fixed Effects

| | Estimate | Std. Error | z value | Pr(>\|z\|) |
|---|---|---|---|---|
| **(Intercept)** | **-2.713** | **1.273** | **-2.130** | **0.033 *** |
| Outcloseness | 1.731 | 1.647 | 1.051 | 0.293 |
| **Valley_Position_UpValley** | **1.915** | **0.436** | **4.393** | **1.12e-05 *** |
| **Predator_Index_Low** | **0.953** | **0.419** | **2.275** | **0.023 *** |
| Pup_Sex_M | 0.036 | 0.215 | 0.167 | 0.868 |
| Litter_Size | -0.018 | 0.064 | -0.282 | 0.778 |
| Network_Size | 0.011 | 0.014 | 0.743 | 0.457 |
| Mother_Age | -0.007 | 0.049 | -0.149 | 0.882 |
| Emergence_Date | 0.021 | 0.019 | 1.107 | 0.268 |
| **Log_Daily_Mass_Gain** | **0.337** | **0.148** | **2.276** | **0.023 *** |
| Outcloseness*Predator_Index_Low | 1.168 | 1.820 | 0.642 | 0.521 |
| **Outcloseness *Valley_Position_UpValley** | **-3.149** | **1.552** | **-2.029** | **0.042 *** |

Outcloseness Random Effects

| Groups Name | Variance | Std. Dev. |
|---|---|---|
| Mother_UID | 0.207 | 0.455 |
| Year | 0.865 | 0.930 |

e. Incloseness Fixed Effects

| | Estimate | Std. Error | z value | Pr(>\|z\|) |
|---|---|---|---|---|
| (Intercept) | -1.856 | 1.277 | -1.454 | 0.146 |
| Incloseness | -0.504 | 2.229 | -0.226 | 0.821 |
| **Valley_Position_UpValley** | **2.052** | **0.428** | **4.791** | **1.66e-06 *** |
| **Predator_Index_Low** | **1.209** | **0.455** | **2.660** | **0.008 *** |
| Pup_Sex_M | 0.002 | 0.002 | 0.008 | 0.994 |
| Litter_Size | -0.029 | 0.062 | -0.465 | 0.642 |
| Network_Size | -0.005 | 0.015 | -0.347 | 0.729 |
| Mother_Age | -0.014 | 0.048 | -0.298 | 0.766 |
| Emergence_Date | 0.006 | 0.018 | 0.351 | 0.726 |
| **Log_Daily_Mass_Gain** | **0.373** | **0.147** | **2.543** | **0.011 *** |
| Incloseness*Predator_Index_Low | 0.095 | 2.122 | 0.045 | 0.964 |
| Incloseness *Valley_Position_UpValley | -2.144 | 1.457 | -1.471 | 0.141 |

Incloseness Random Effects

| Groups Name | Variance | Std. Dev. |
|---|---|---|
| Mother_UID | 0.151 | 0.388 |
| Year | 0.719 | 0.848 |

f. Local Clustering Fixed Effects

| | Estimate | Std. Error | z value | Pr(>\|z\|) |
|---|---|---|---|---|
| (Intercept) | -1.274 | 1.377 | -0.926 | 0.355 |
| Local_Clustering | 0.457 | 0.841 | 0.543 | 0.587 |
| **Valley_Position_UpValley** | **1.270** | **0.444** | **2.860** | **0.004 *** |
| Predator_Index_Low | 0.635 | 0.435 | 1.460 | 0.144 |
| Pup_Sex_M | 0.040 | 0.236 | 0.172 | 0.864 |
| Litter_Size | -0.001 | 0.067 | -0.019 | 0.985 |
| Network_Size | -0.004 | 0.018 | -0.228 | 0.820 |
| Mother_Age | 2.37e-04 | 0.053 | 0.004 | 0.996 |
| Emergence_Date | 0.007 | 0.020 | 0.359 | 0.720 |
| Log_Daily_Mass_Gain | 0.193 | 0.161 | 1.199 | 0.231 |
| Local_Clustering *Predator_Index_Low | 1.670 | 1.104 | 1.513 | 0.130 |
| Local_Clustering *Valley_Position_UpValley | 1.034 | 1.126 | 0.918 | 0.359 |

Local Clustering Random Effects

| Groups Name | Variance | Std. Dev. |
|---|---|---|
| Mother_UID | 0.314 | 0.561 |
| Year | 0.812 | 0.901 |

g. Negative Average Shortest Path Length Fixed Effects

| | Estimate | Std. Error | z value | Pr(>\|z\|) |
|---|---|---|---|---|
| (Intercept) | 0.446 | 1.492 | 0.299 | 0.765 |
| **Neg_Average_Shortest_Path_Length** | **1.362** | **0.534** | **2.549** | **0.011 *** |
| **Valley_Position_UpValley** | **-2.653** | **1.213** | **-2.187** | **0.029 *** |
| Predator_Index_Low | 1.251 | 1.198 | 1.045 | 0.296 |
| Pup_Sex_M | 0.020 | 0.215 | 0.095 | 0.924 |
| Litter_Size | -0.017 | 0.061 | -0.283 | 0.777 |
| Network_Size | 0.017 | 0.017 | 0.989 | 0.323 |
| Mother_Age | -0.010 | 0.048 | -0.223 | 0.824 |
| Emergence_Date | 0.013 | 0.018 | 0.737 | 0.461 |
| **Log_Daily_Mass_Gain** | **0.313** | **0.142** | **2.194** | **0.028 *** |
| Neg_Average_Shortest_Path_Length *Predator_Index_Low | -0.005 | 0.570 | -0.010 | 0.992 |
| **Neg_Average_Shortest_Path_Length *Valley_Position_UpValley** | **-1.917** | **0.558** | **-3.434** | **0.001 *** |

Negative Average Shortest Path Length Random Effects

| Groups Name | Variance | Std. Dev. |
|---|---|---|
| Mother_UID | 0.144 | 0.379 |
| Year | 0.827 | 0.909 |

h. Eigenvector Centrality Fixed Effects

| | Estimate | Std. Error | z value | Pr(>\|z\|) |
|---|---|---|---|---|
| (Intercept) | -2.375 | 1.222 | -1.944 | 0.052 |
| Eigenvector_Centrality | 0.241 | 0.674 | 0.358 | 0.720 |
| **Valley_Position_UpValley** | **1.827** | **0.383** | **4.772** | **1.83e-06 *** |
| **Predator_Index_Low** | **0.859** | **0.361** | **2.380** | **0.017 *** |
| Pup_Sex_M | 0.026 | 0.214 | 0.122 | 0.903 |
| Litter_Size | -0.021 | 0.063 | -0.339 | 0.735 |
| Network_Size | 0.010 | 0.014 | 0.725 | 0.468 |
| Mother_Age | -0.012 | 0.049 | -0.254 | 0.800 |
| Emergence_Date | 0.018 | 0.019 | 0.949 | 0.343 |
| **Log_Daily_Mass_Gain** | **0.355** | **0.147** | **2.421** | **0.015 *** |
| Eigenvector_Centrality *Predator_Index_Low | 0.797 | 0.871 | 0.915 | 0.360 |
| Eigenvector_Centrality *Valley_Position_UpValley | -1.528 | 0.801 | -1.907 | 0.057 |

Eigenvector Centrality Random Effects

| Groups Name | Variance | Std. Dev. |
|---|---|---|
| Mother_UID | 0.194 | 0.440 |
| Year | 0.821 | 0.906 |

i. Outstrength Fixed Effects

| | Estimate | Std. Error | z value | Pr(>\|z\|) |
|---|---|---|---|---|
| (Intercept) | -2.340 | 1.225 | -1.911 | 0.056 |
| Outstrength | 0.031 | 0.029 | 1.059 | 0.290 |
| **Valley_Position_UpValley** | **1.929** | **0.351** | **5.501** | **3.78e-08 *** |
| **Predator_Index_Low** | **1.230** | **0.331** | **3.712** | **2.05e-04 *** |
| Pup_Sex_M | 0.010 | 0.214 | 0.046 | 0.963 |
| Litter_Size | -0.023 | 0.063 | -0.361 | 0.718 |
| Network_Size | 0.002 | 0.014 | 0.172 | 0.863 |
| Mother_Age | -0.022 | 0.050 | -0.432 | 0.666 |
| Emergence_Date | 0.009 | 0.018 | 0.502 | 0.616 |
| **Log_Daily_Mass_Gain** | **0.381** | **0.147** | **2.603** | **0.009 *** |
| Outstrength*Predator_Index_Low | -0.011 | 0.030 | -0.376 | 0.707 |
| Outstrength *Valley_Position_UpValley | -0.048 | 0.025 | -1.925 | 0.054 |

Outstrength Random Effects

| Groups Name | Variance | Std. Dev. |
|---|---|---|
| Mother_UID | 0.189 | 0.434 |
| Year | 0.695 | 0.834 |

j. Instrength Fixed Effects

| | Estimate | Std. Error | z value | Pr(>\|z\|) |
|---|---|---|---|---|
| (Intercept) | -1.872 | 1.280 | -1.463 | 0.144 |
| Instrength | -0.025 | 0.019 | -1.306 | 0.192 |
| **Valley_Position_UpValley** | **1.746** | **0.351** | **4.970** | **6.69e-07 *** |
| **Predator_Index_Low** | **0.911** | **0.335** | **2.720** | **0.006 *** |
| Pup_Sex_M | 0.019 | 0.215 | 0.086 | 0.931 |
| Litter_Size | -0.030 | 0.064 | -0.470 | 0.638 |
| Network_Size | 0.012 | 0.014 | 0.851 | 0.395 |
| Mother_Age | -0.018 | 0.049 | -0.374 | 0.708 |
| Emergence_Date | 0.007 | 0.018 | 0.406 | 0.685 |
| **Log_Daily_Mass_Gain** | **0.348** | **0.151** | **2.299** | **0.021 *** |
| Instrength*Predator_Index_Low | 0.026 | 0.020 | 1.267 | 0.205 |
| Instrength *Valley_Position_UpValley | -0.017 | 0.016 | -1.065 | 0.287 |

Instrength Random Effects

| Groups Name | Variance | Std. Dev. |
|---|---|---|
| Mother_UID | 0.180 | 0.424 |
| Year | 0.806 | 0.898 |

**Supplementary Table 4. Agonistic Yearly Survival Models**

Tables show the complete GLMM fixed and random effects outputs for the yearly survival models based on agonistic social measures. Social measures include **a**. indegree, **b.** outdegree, **c.** betweenness, **d.** outcloseness, **e.** incloseness, **f.** local clustering, **g.** average shortest path length, **h.** eigenvector centrality, **i.** outstrength, **j.** instrength. Significant variables are bolded and marked with an asterisk.

a. Indegree Fixed Effects

| | Estimate | Std. Error | z value | Pr(>\|z\|) |
|---|---|---|---|---|
| (Intercept) | -0.580 | 0.749 | -0.775 | 0.438 |
| Indegree | 0.148 | 1.974 | 0.075 | 0.940 |
| **Valley_Position_UpValley** | **0.875** | **0.372** | **2.353** | **0.019 *** |
| **Predator_Index_Low** | **1.241** | **0.379** | **3.275** | **0.001 *** |
| Pup_Sex_M | 0.081 | 0.183 | 0.440 | 0.660 |
| Litter_Size | -0.062 | 0.056 | -1.108 | 0.268 |
| Network_Size | 0.018 | 0.021 | 0.875 | 0.381 |
| Mother_Age | 0.027 | 0.043 | 0.635 | 0.526 |
| Emergence_Date | -0.025 | 0.015 | -1.619 | 0.105 |
| Indegree*Predator_Index_Low | -0.997 | 2.011 | -0.496 | 0.620 |
| Indegree* Valley_Position_UpValley | 0.588 | 1.568 | 0.375 | 0.707 |

Indegree Random Effects

| Groups Name | Variance | Std. Dev. |
|---|---|---|
| Mother_UID | 0.0867 | 0.294 |
| Year | 1.527 | 1.236 |

b. Outdegree Fixed Effects

| | Estimate | Std. Error | z value | Pr(>\|z\|) |
|---|---|---|---|---|
| **(Intercept)** | **-8.266** | **1.533** | **-5.390** | **7.04e-08 *** |
| Outdegree | 2.357 | 2.154 | 1.095 | 0.274 |
| **Valley_Position_UpValley** | **1.469** | **0.364** | **4.034** | **5.49e-05 *** |
| **Predator_Index_Low** | **1.393** | **0.389** | **3.578** | **3.46e-04 *** |
| Pup_Sex_M | -0.195 | 0.195 | -0.999 | 0.318 |
| Litter_Size | -0.006 | 0.058 | -0.102 | 0.919 |
| Network_Size | 0.039 | 0.021 | 1.898 | 0.058 |
| Mother_Age | 0.009 | 0.044 | 0.198 | 0.843 |
| **Emergence_Date** | **0.047** | **0.020** | **2.417** | **0.016 *** |
| **Log_August_Mass** | **0.949** | **0.176** | **5.397** | **6.77e-08 *** |
| Outdegree*Predator_Index_Low | -0.119 | 2.114 | -0.056 | 0.955 |
| Outdegree *Valley_Position_UpValley | -2.547 | 1.718 | -1.483 | 0.138 |

Outdegree Random Effects

| Groups Name | Variance | Std. Dev. |
|---|---|---|
| Mother_UID | 0.050 | 0.223 |
| Year | 1.685 | 1.298 |

c. Betweenness Fixed Effects

| | Estimate | Std. Error | z value | Pr(>\|z\|) |
|---|---|---|---|---|
| (Intercept) | -0.698 | 0.643 | -1.085 | 0.278 |
| Betweenness | -3.395 | 2.639 | -1.286 | 0.198 |
| **Valley_Position_UpValley** | **0.852** | **0.261** | **3.263** | **0.001 *** |
| **Predator_Index_Low** | **1.004** | **0.267** | **3.764** | **1.67e-04 *** |
| Pup_Sex_M | 0.088 | 0.183 | 0.484 | 0.629 |
| Litter_Size | -0.034 | 0.055 | -0.610 | 0.542 |
| Network_Size | 0.026 | 0.019 | 1.394 | 0.163 |
| Mother_Age | 0.029 | 0.041 | 0.703 | 0.482 |
| Emergence_Date | -0.029 | 0.015 | -1.932 | 0.053 |
| Betweenness*Predator_Index_Low | 3.380 | 2.797 | 1.208 | 0.227 |
| Betweenness *Valley_Position_UpValley | 4.237 | 2.503 | 1.693 | 0.090 |

Betweenness Random Effects

| Groups Name | Variance | Std. Dev. |
|---|---|---|
| Mother_UID | 0.048 | 0.219 |
| Year | 1.533 | 1.238 |

d. Outcloseness Fixed Effects

| | Estimate | Std. Error | z value | Pr(>\|z\|) |
|---|---|---|---|---|
| **(Intercept)** | **-8.295** | **1.588** | **-5.223** | **1.76e-07 *** |
| Outcloseness | 0.365 | 1.842 | 0.198 | 0.843 |
| **Valley_Position_UpValley** | **1.304** | **0.376** | **3.464** | **5.32e-04 *** |
| **Predator_Index_Low** | **1.085** | **0.407** | **2.663** | **0.008 *** |
| Pup_Sex_M | -0.195 | 0.196 | -0.998 | 0.318 |
| Litter_Size | -0.001 | 0.059 | -0.023 | 0.982 |
| Network_Size | 0.043 | 0.022 | 1.928 | 0.054 |
| Mother_Age | 0.005 | 0.044 | 0.122 | 0.903 |
| **Emergence_Date** | **0.054** | **0.020** | **2.638** | **0.008 *** |
| **Log_August_Mass** | **0.970** | **0.178** | **5.440** | **5.32e-08 *** |
| Outcloseness*Predator_Index_Low | 1.613 | 1.841 | 0.876 | 0.381 |
| Outcloseness *Valley_Position_UpValley | -1.515 | 1.271 | -1.192 | 0.233 |

Outcloseness Random Effects

| Groups Name | Variance | Std. Dev. |
|---|---|---|
| Mother_UID | 0.056 | 0.238 |
| Year | 1.861 | 1.364 |

e. Incloseness Fixed Effects

| | Estimate | Std. Error | z value | Pr(>\|z\|) |
|---|---|---|---|---|
| (Intercept) | -0.187 | 0.796 | -0.235 | 0.814 |
| Incloseness | -2.191 | 2.336 | -0.938 | 0.348 |
| **Valley_Position_UpValley** | **1.257** | **0.361** | **3.485** | **4.91e-04 *** |
| Predator_Index_Low | 0.668 | 0.424 | 1.577 | 0.115 |
| Pup_Sex_M | 0.078 | 0.183 | 0.428 | 0.669 |
| Litter_Size | -0.060 | 0.055 | -1.088 | 0.276 |
| Network_Size | 0.010 | 0.022 | 0.483 | 0.629 |
| Mother_Age | 0.034 | 0.042 | 0.797 | 0.425 |
| Emergence_Date | -0.027 | 0.015 | -1.764 | 0.078 |
| Incloseness*Predator_Index_Low | 2.942 | 2.181 | 1.349 | 0.177 |
| Incloseness *Valley_Position_UpValley | -1.347 | 1.301 | -1.035 | 0.301 |

Incloseness Random Effects

| Groups Name | Variance | Std. Dev. |
|---|---|---|
| Mother_UID | 0.061 | 0.247 |
| Year | 1.566 | 1.251 |

f. Local Clustering Fixed Effects

| | Estimate | Std. Error | z value | Pr(>\|z\|) |
|---|---|---|---|---|
| **(Intercept)** | **-8.067** | **1.561** | **-5.168** | **2.37e-07 *** |
| Local_Clustering | 0.928 | 0.789 | 1.176 | 0.239 |
| **Valley_Position_UpValley** | **1.328** | **0.373** | **3.559** | **3.72e-04 *** |
| **Predator_Index_Low** | **1.318** | **0.395** | **3.333** | **0.001 *** |
| Pup_Sex_M | -0.283 | 0.207 | -1.371 | 0.170 |
| Litter_Size | 0.010 | 0.058 | 0.170 | 0.865 |
| Network_Size | 0.046 | 0.024 | 1.891 | 0.059 |
| Mother_Age | 0.011 | 0.045 | 0.238 | 0.812 |
| **Emergence_Date** | **0.044** | **0.021** | **2.132** | **0.033 *** |
| **Log_August_Mass** | **0.935** | **0.183** | **5.103** | **3.34e-07 *** |
| Local_Clustering *Predator_Index_Low | -0.140 | 0.879 | -0.159 | 0.874 |
| Local_Clustering *Valley_Position_UpValley | -0.911 | 0.834 | -1.093 | 0.274 |

Local Clustering Random Effects

| Groups Name | Variance | Std. Dev. |
|---|---|---|
| Mother_UID | 0.089 | 0.299 |
| Year | 1.720 | 1.311 |

g. Negative Average Shortest Path Length Fixed Effects

| | Estimate | Std. Error | z value | Pr(>\|z\|) |
|---|---|---|---|---|
| (Intercept) | 2.163 | 1.143 | 1.892 | 0.059 |
| **Neg_Average_Shortest_Path_Length** | **1.633** | **0.537** | **3.042** | **0.002 *** |
| Valley_Position_UpValley | -1.907 | 1.056 | -1.806 | 0.071 |
| Predator_Index_Low | 0.807 | 1.018 | 0.793 | 0.428 |
| Pup_Sex_M | 0.058 | 0.184 | 0.316 | 0.752 |
| Litter_Size | -0.070 | 0.055 | -1.266 | 0.206 |
| Network_Size | 0.053 | 0.027 | 1.957 | 0.050 |
| Mother_Age | 0.028 | 0.042 | 0.658 | 0.511 |
| Emergence_Date | -0.022 | 0.015 | -1.480 | 0.139 |
| Neg_Average_Shortest_Path_Length *Predator_Index_Low | -0.169 | 0.477 | -0.354 | 0.724 |
| **Neg_Average_Shortest_Path_Length *Valley_Position_UpValley** | **-1.468** | **0.498** | **-2.946** | **0.003 *** |

Negative Average Shortest Path Length Random Effects

| Groups Name | Variance | Std. Dev. |
|---|---|---|
| Mother_UID | 0.062 | 0.250 |
| Year | 1.470 | 1.212 |

h. Eigenvector Centrality Fixed Effects

| | Estimate | Std. Error | z value | Pr(>\|z\|) |
|---|---|---|---|---|
| **(Intercept)** | **-7.641** | **1.449** | **-5.272** | **1.35e-07 *** |
| Eigenvector_Centrality | -0.077 | 0.714 | -0.108 | 0.914 |
| **Valley_Position_UpValley** | **0.951** | **0.330** | **2.881** | **0.004 *** |
| **Predator_Index_Low** | **1.419** | **0.339** | **4.187** | **2.83e-05 *** |
| Pup_Sex_M | -0.190 | 0.194 | -0.982 | 0.326 |
| Litter_Size | -0.002 | 0.056 | -0.032 | 0.975 |
| Group_Size | 0.037 | 0.020 | 1.860 | 0.063 |
| Mother_Age | 0.008 | 0.043 | 0.190 | 0.849 |
| **Emergence_Date** | **0.043** | **0.019** | **2.196** | **0.028 *** |
| **Log_August_Mass** | **0.927** | **0.172** | **5.377** | **7.56e-08 *** |
| Eigenvector_Centrality *Predator_Index_Low | -0.329 | 0.801 | -0.411 | 0.681 |
| Eigenvector_Centrality *Valley_Position_UpValley | 0.495 | 0.733 | 0.676 | 0.499 |

Eigenvector Centrality Random Effects

| Groups Name | Variance | Std. Dev. |
|---|---|---|
| Mother_UID | 0.016 | 0.128 |
| Year | 1.791 | 1.338 |

i. Outstrength Fixed Effects

| | Estimate | Std. Error | z value | Pr(>\|z\|) |
|---|---|---|---|---|
| (Intercept) | -0.511 | 0.670 | -0.763 | 0.446 |
| Outstrength | 0.008 | 0.027 | 0.296 | 0.767 |
| **Valley_Position_UpValley** | **1.218** | **0.289** | **4.211** | **2.54e-05 *** |
| **Predator_Index_Low** | **1.155** | **0.286** | **4.033** | **5.50e-05 *** |
| Pup_Sex_M | 0.089 | 0.183 | 0.485 | 0.628 |
| Litter_Size | -0.067 | 0.055 | -1.204 | 0.228 |
| Network_Size | 0.018 | 0.020 | 0.911 | 0.362 |
| Mother_Age | 0.018 | 0.043 | 0.426 | 0.670 |
| Emergence_Date | -0.029 | 0.015 | -1.882 | 0.060 |
| Outstrength*Predator_Index_Low | -0.003 | 0.027 | -0.107 | 0.915 |
| Outstrength *Valley_Position_UpValley | -0.022 | 0.017 | -1.317 | 0.188 |

Outstrength Random Effects

| Groups Name | Variance | Std. Dev. |
|---|---|---|
| Mother_UID | 0.060 | 0.245 |
| Year | 1.443 | 1.201 |

j. Instrength Fixed Effects

| | Estimate | Std. Error | z value | Pr(>\|z\|) |
|---|---|---|---|---|
| (Intercept) | 0.049 | 0.656 | 0.075 | 0.940 |
| Instrength | -0.037 | 0.0191 | -1.942 | 0.052 |
| **Valley_Position_UpValley** | **0.925** | **0.290** | **3.190** | **0.001 *** |
| **Predator_Index_Low** | **1.052** | **0.283** | **3.714** | **2.04e-04 *** |
| Pup_Sex_M | 0.061 | 0.184 | 0.332 | 0.740 |
| Litter_Size | -0.091 | 0.055 | -1.664 | 0.096 |
| Network_Size | 0.021 | 0.019 | 1.149 | 0.250 |
| Mother_Age | 0.024 | 0.041 | 0.589 | 0.556 |
| **Emergence_Date** | **-0.032** | **0.015** | **-2.126** | **0.033 *** |
| Instrength*Predator_Index_Low | 0.009 | 0.020 | 0.474 | 0.636 |
| Instrength *Valley_Position_UpValley | 0.014 | 0.014 | 0.987 | 0.324 |

Instrength Random Effects

| Groups Name | Variance | Std. Dev. |
|---|---|---|
| Mother_UID | 0.014 | 0.118 |
| Year | 1.428 | 1.195 |
